# BCOR mutations establish a persistent culture-adaptive state in human induced pluripotent stem cells

**DOI:** 10.64898/2026.09.04.739550

**Authors:** Cherif Badja, Soraya Boushaki, Yogesh Kumar, Theodoros I. Roumeliotis, Annabel J. Curle, Angelica Elizabeth Guiloff, Andrea Degasperi, Sophie Momen, Florian Robert, Yasin Memari, Scott Shooter, Zuza Kozik, Salome Jingchen Zhao, Ping Jing Toong, India Baker, Brian Hendrich, Roger Alistair Barker, Jyoti Choudhary, Gene Ching Cheik Koh, Serena Nik-Zainal

## Abstract

Human induced pluripotent stem cells (hiPSCs) are widely used for disease modelling and regenerative medicine, yet their utility depends on maintaining molecular integrity during long-term culture. BCOR (BCL6 Co-Repressor) mutations are among the most recurrent culture-acquired alterations in hiPSCs, but their functional consequences remain poorly understood. Here, we show that hiPSC *BCOR* mutations are predominantly truncating indels enriched in exon 7, defining a mutational landscape distinct from that observed in cancer. Multi-omics profiling reveals that BCOR loss drives widespread chromatin, transcriptomic and proteomic remodelling, with coordinated activation of developmental, pluripotency-associated and mitochondrial metabolism programmes. To facilitate routine surveillance, we develop a cost-effective TaqMan qPCR assay that accurately identifies BCOR-mutant hiPSCs across independent cell lines. Finally, we demonstrate that correction of BCOR mutation by CRISPR–Cas9 only partially restores the wildtype molecular state, highlighting the importance of early detection and monitoring of adaptive mutations in hiPSC cultures.

## Introduction

Twenty years have passed since the first demonstration of induced pluripotency in mouse cells(Takahashi and Yamanaka, 2006; Yamanaka, 2026). Induced pluripotent stem cells (iPSCs) have moved from a laboratory curiosity to a foundational technology underpinning disease modelling(Badja et al., 2024; Mazaleyrat et al., 2020), drug discovery, regenerative therapy and rejuvenation research(Mandai et al., 2017; Sawamoto et al., 2025; Schweitzer et al., 2020; Yamanaka, 2026). Patient-derived hiPSCs offer a uniquely powerful experimental system that can capture the full genetic background of an individual, self-renew indefinitely in culture (in theory), and differentiate into derivatives of all three germ layers. They circumvent the ethical constraints associated with human embryonic stem cells, enable autologous and HLA-matched cell therapies that minimise immune rejection, and have already entered the clinic in trials for Parkinson’s disease(Sawamoto *et al*., 2025; Schweitzer *et al*., 2020) and age-related macular degeneration(Mandai *et al*., 2017). Beyond the clinic, hiPSCs have underpinned large-scale pharmacogenomic screens(Roberts et al., 2019), *in vitro* models of human development(Shahbazi et al., 2019), mutagen and DNA-repair screens that decode the origins of cancer mutational signatures(Kucab et al., 2019; Zou et al., 2021), and an expanding catalogue of organoid systems that recapitulate tissue architecture and pathology in a dish(Lancaster et al., 2013).

The realisation of these applications has rested as much on collective community infrastructure as on individual scientific breakthroughs. Coordinated efforts by HipSci(Kilpinen et al., 2017; Streeter et al., 2017), the Allen Institute(Roberts *et al*., 2019), EBiSC(De Sousa et al., 2017; Steeg et al., 2020), WiCell(Kim et al., 2017), CIRM(Lin et al., 2020), the Human Pluripotent Stem Cell Registry(Seltmann et al., 2016) and many others have generated thousands of well-characterised lines, established standards for derivation, banking, identity testing and karyotyping, and made hiPSCs widely accessible. Hand in hand with this expansion, the community has progressively confronted the practical challenges of working with pluripotent cells, including tumorigenicity(Yasuda et al., 2018), immunogenicity(Zhao et al., 2011) and line-to-line heterogeneity(Yamanaka, 2020), and developed safer reprogramming strategies, footprint-free vectors, defined culture conditions, and increasingly stringent genomic surveillance. This collective effort to ensure the safety, reproducibility and biological fidelity of hiPSC models is now a central pillar of the field and is essential if the technology is to deliver on its therapeutic promise in the clinic.

A persistent challenge in this regard is the appearance of culture-acquired genetic alterations that escape standard quality-control checks(Benvenisty et al., 2025). hiPSCs are subject to selective pressure during reprogramming, single-cell subcloning and prolonged passaging, with variants outcompeting wildtype through mechanisms such as YAP-mediated cell competition(Price et al., 2021) and culture condition-dependent selection that, for example, favours chromosome 1q gains in feeder-free regimens(Stavish et al., 2024). Recurrent mutations have been documented in genes including *TP53*(Gore et al., 2011; Merkle et al., 2017), *DNMT3A*, *TET2* and recently, *BCOR*(Puigdevall et al., 2023; Rouhani et al., 2022) was found to be one of the most frequently acquired mutations in blood-derived hiPSCs in particular.

*BCOR* is a core component of the non-canonical Polycomb Repressive Complex 1.1 (ncPRC1.1). It cooperates with *KDM2B* and *RING1A/B* to recognise unmethylated CpG islands and repress genes governing differentiation, metabolism and cell-cycle control independently of the H3K27me3 mark deposited by PRC2(Farcas et al., 2012; Gao et al., 2012; Zhu et al., 2022). Germline *BCOR* mutations are causally linked to oculofaciocardiodental syndrome, while somatic *BCOR* mutations are reported drivers in acute myeloid leukaemia, CNS high-grade neuroepithelial tumour and endometrial carcinoma(Honda et al., 2021; Momeni-Boroujeni et al., 2023; Ng et al., 2004; Sturm et al., 2016). In hiPSCs, *BCOR* mutations are positively selected and display altered neural differentiation(Puigdevall *et al*., 2023; Rouhani *et al*., 2022). Despite this clinical and experimental significance, the molecular consequences of *BCOR* mutations in hiPSCs have not been extensively characterised, nor do we fully understand whether they can be prevented or reversed. Equally, the field lacks an accessible, low-cost screening assay that allows routine surveillance of *BCOR* status during long-term culture and clonal expansion.

In this study, we address these outstanding questions by combining whole-genome sequencing, ChIP-seq, transcriptomics and proteomics to define the orchestrated molecular consequences of BCOR loss in hiPSCs across multiple regulatory layers. We then translate these findings into a practical cDNA-based TaqMan assay and validate its performance in independent hiPSC lines from external laboratories. Finally, we ask whether precise CRISPR-Cas9 correction of a recurrent exon 7 *BCOR* indel is sufficient to revert the BCOR mutant state and restore the molecular features of wild-type cells. Our findings provide new insights into one of the most recurrently selected mutations in hiPSC culture and establish an accessible tool for routine surveillance of BCOR status, helping to safeguard the integrity of hiPSC models as their technological applications continue to expand.

## Results

### BCOR mutations in hiPSCs are mainly truncating loss-of-function indel mutations enriched in exon7

We first examined the distribution of reported BCOR mutations in hiPSCs derived from blood and from skin fibroblasts(Puigdevall *et al*., 2023; Rouhani *et al*., 2022). Of 55 reported BCOR mutations(Puigdevall *et al*., 2023; Rouhani *et al*., 2022), 53 (96.36%) were truncating variants, of which 79.25% (42/53) were frameshifting indels and 20.75% (11/53) were nonsense substitutions. Only two mutations were missense mutations (*p*.P1128S in exon 7 and *p*.R1375W in exon 9).

Comparison of BCOR mutation distributions across hiPSCs, AMLs and non-AML cancers revealed marked differences (Figure 1A-H). In hiPSCs, mutations were strongly enriched in exon 7, which accounted for 47.3% of all reported variants and was significantly overrepresented relative to both AML (Fisher’s test, *P*-value: 1.49E-07) (Figure 1A, 1B), and non-AML cancers (Fisher’s test, *P*-value:1.21E-20) (Figure 1A, 1C). By contrast, AML-associated BCOR mutations were concentrated in exons 4 (52.3%; Fisher’s test, *P*-value: 0.013) and 9 (18.2%; Fisher’s test, *P*-value: 0.021) whereas non-AML cancers showed enrichment in exons 10 (34.6%; Fisher’s test, *P*-value: 1.38E-06) and 4 (32.5%; Fisher’s test, *P*-value: 0.45) (Figure 1D, 1E, 1F). In addition, non-AML cancers exhibited a marked enrichment for the N1459S/T mutation in exon 10, with nearly 25% of the cohort carrying this specific missense variant (176/709), a pattern not observed in either AMLs or hiPSCs (Figure 1C). These observations suggest that selective pressure acting on BCOR are highly context-dependent, favouring distinct mutational spectra in hiPSCs, AML and other cancer types.

**Figure 1.**
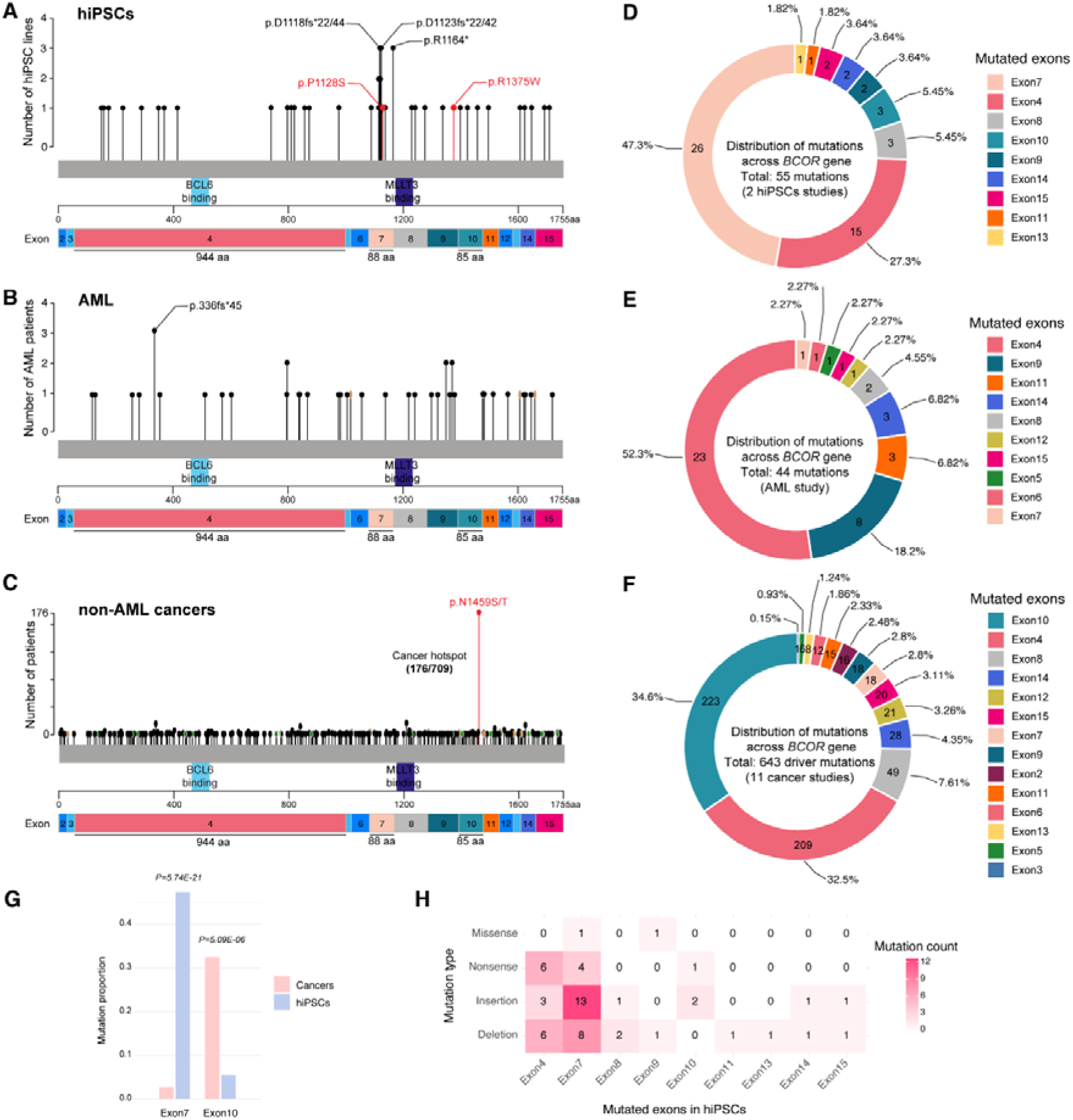
**BCOR mutations in hiPSCs are enriched in exon 7 and display a distinct mutational spectrum from cancer-associated BCOR mutations. A–C**, Distribution of BCOR mutations across the BCOR coding sequence in hiPSCs (*n* = 55 mutations from two independent studies), AML (*n* = 44 mutations – www.cbioportal.org) and non-AML cancers (*n* = 643 driver mutations from 11 studies –cBioPortal). Recurrent mutations and hotspot residues are indicated. **D–F**, Exon-level distribution of BCOR mutations in hiPSCs, AML and non-AML cancers. **G**, Comparison of mutation frequencies in exon 7 and exon 10 between hiPSC and cancer datasets. Exon 7 mutations were significantly enriched in hiPSCs relative to cancers (two-sided Fisher’s exact test, *P* = 5.74 × 10⁻²¹), whereas exon 10 mutations were significantly enriched in cancer (*P* = 5.09 × 10⁻). **H**, Distribution of mutation classes across BCOR exons in hiPSCs, showing the predominance of truncating indels within exon.

To investigate why exon 7 is preferentially mutated in hiPSCs, we examined the DNA sequence context of the reported variants. Half of the exon 7 mutations were insertions (13/26), eight were deletions, four were nonsense mutations and one was a missense mutation (Figure 1H). Intriguingly, a recurrent 17bp sequence motif “AGGCCACCTGGTCTGCG” located in the centre of exon 7 was repeatedly inserted or deleted across independent lines. Furthermore, most BCOR mutations (53/55) occurred either at G/C nucleotides or immediately adjacent to them (Supp Figure 1A).

Motif analysis revealed enrichment of BCOR mutations at computationally predicted non-canonical G-quadruplex sequences (G4Q)(Kikin et al., 2006), identified using QGRS Mapper with a conservative G-score threshold of greater than 30 (Supp Figure 1B). Predicted G4Q motifs were distributed throughout the BCOR gene body but were particularly enriched in exons 4, 7, 8, 9, 10 (Supp Figure 1B). Because G-quadruplex structures are located predominantly at enhancers and promoters(Su et al., 2025) and have been associated with genomic instability and mutagenesis(Su *et al*., 2025; Zhuk et al., 2024), we next examined the distribution of DNase I Hypersensitive Sites (DHSs). Notably, exon 7 was the only BCOR exon containing a DHS detected in pluripotent stem cells (H1, H9, iPSC) that overlapped with predicted G4Q sequences. Our observation raises the possibility that the recurrent enrichment of BCOR mutations in exon 7 may be influenced by local sequence and chromatin features in hiPSCs. Direct testing of this hypothesis will require future functional studies.

### BCOR mutant hiPSCs show widespread chromatin dysregulation and enhanced resilience to DNA damage

BCOR loss-of-function mutations have previously been shown to alter the transcriptional programme of hiPSCs, impairing neural differentiation(Puigdevall *et al*., 2023; Rouhani *et al*., 2022; Wang et al., 2018). However, their impact on the epigenome has not been investigated. Given the role of the ncPRC1.1 complex in chromatin-mediated gene regulation, we examined the distribution of active (H3K27ac, H3K4me3) and repressive histone marks (H3K27me3) in BCOR-mutant and wildtype (BCOR-Wt) B-hiPSCs.

ChIP-seq analysis revealed a widespread epigenetic dysregulation in BCOR-mutant cells, affecting all three chromatin marks examined (Figure 2A-2C, Supp Figure 2A-C). Gene Ontology analysis of genes associated with regions depleted of H3K27me3 (Figure 2D) or enriched for H3K27ac (Figure 2E) and H3K4me3 (Figure 2F) identified significant enrichment of developmental and differentiation pathways, consistent with an important role for *BCOR* in maintaining pluripotent cell identity and regulating lineage commitment. As expected, loss of H3K27me3 was associated with increased deposition of H3K4me3 (*R*= −0.507, n= 108 genes) (Figure 2G), and H3K27ac (*R*= −0.727, n=97 genes) (Figure 2H) while gains in H3K27ac and H3K4me3 were positively correlated (*R*= 0.688, n=38 genes) (Figure 2I). Overall, these findings indicate that BCOR loss drives global (rather than focal) redistribution and dysregulation of both active and repressive chromatin marks across the genome.

**Figure 2.**
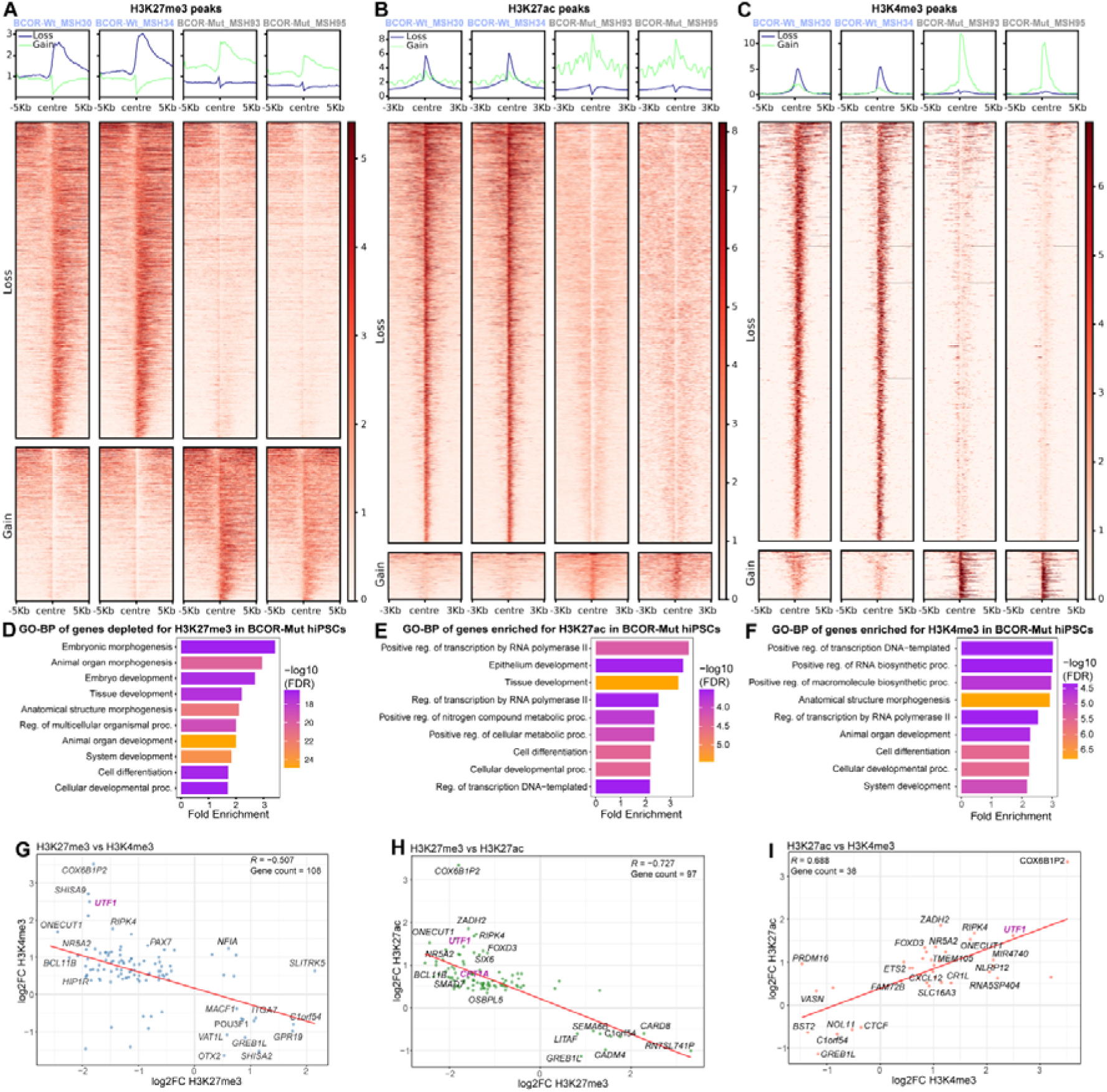
**BCOR mutations reshape active and repressive chromatin landscapes in human iPSCs. A–C**, Differential enrichment of H3K27me3, H3K27ac and H3K4me3 peaks between BCOR-wildtype and BCOR-mutant hiPSCs. Representative data are shown for two independent BCOR-wildtype lines (MSH30 and MSH34) and two independent BCOR-mutant lines (MSH93 and MSH95). **D–F**, Gene Ontology biological process (GO-BP) enrichment analyses of genes associated with differential chromatin regions. **G–I**, Correlation analyses of genes showing coordinated changes across chromatin states. Changes in H3K27me3 were inversely correlated with H3K4me3 (*R* = −0.507, *n* = 108 genes; **G**) and H3K27ac (*R* = −0.727, *n* = 97 genes; **H**), whereas H3K27ac and H3K4me3 changes were positively correlated (*R* = 0.688, *n* = 38 genes; **I**). Selected genes involved in pluripotency, developmental regulation and cellular metabolism are highlighted.

Beyond developmental pathways, a subset of loci gaining active chromatin marks were associated with cellular growth, metabolic adaptation and stress-response programmes. These included *UTF1*(Jerber et al., 2021; Lou et al., 2023; Mouallif et al., 2014; Okuda et al., 1998), *CPT1A*(Jariwala et al., 2021; Xiong et al., 2020), NR5A2(Gu et al., 2005; Heng et al., 2010; Ye et al., 2019) and *RIPK4*(Liu et al., 2018; Liu et al., 2021), genes previously implicated in pluripotency, proliferation, metabolic reprogramming, therapy resistance and tumour progression (Figure 2G–I, Supplementary Figure 2D–F). Collectively, these observations suggested that BCOR mutations may promote cellular fitness and adaptation to stress.

To test this hypothesis, we used a previously established hiPSC models of Xeroderma Pigmentosum (XP)(Badja *et al*., 2024), which carry mutations altering the Nucleotide Excision Repair (NER) pathway. Consistent with their DNA repair deficiency, NER-deficient BCOR-wildtype B-hiPSCs (XP^Mut^/BCOR^Wt^) exhibited extensive cell death and loss of colony-forming capacity following a single low dose of UVC irradiation (0.75 J/m²) (Supplementary Figure 2H). In contrast, NER-deficient B-hiPSCs carrying a BCOR mutation (XP^Mut^/BCORMut) showed markedly improved survival under the same conditions (Supplementary Figure 2I). Together, these findings suggest that BCOR mutations enhance tolerance to DNA damage and may contribute to the selective advantage of BCOR-mutant hiPSCs during long-term culture.

### BCOR-mutant transcriptional and proteomic states mirror chromatin dysregulation and reveal metabolic rewiring

To determine whether the chromatin alterations observed in BCOR-mutant (BCOR-Mut) B-hiPSCs were reflected at the transcriptional and protein levels, we performed RNA-seq and quantitative proteomic profiling.

Principal component analysis of the transcriptomic data revealed clear segregation of samples according to BCOR status (Figure 3A), indicating that BCOR mutations are a major determinant of transcriptional variance in hiPSCs. Differential expression analysis identified 10,347 genes (padj <0.05), including 4,530 upregulated and 5,817 downregulated genes in BCOR-mutant lines (Figure 3B). Pathway enrichment analysis (KEGG Gene Ontology analysis (ShinyGo 0.85.1)) revealed significant upregulation of oxidative phosphorylation, reactive oxygen species-associated pathways, metabolism and cancer-related pathways, whereas neurodevelopmental and neural signalling pathways related to glutamatergic activity, neuroactive ligand-receptor interactions, and calcium signalling were significantly downregulated (Supp Figure 3A, B). Consistent with these findings, genes implicated in pluripotency and developmental patterning including but not limited to *UTF1*, *IRX4*, *FOXC1*, *HOXB13*, *PRAC1*, *PITX2*, *ZIC1*, *NODAL were upregulated*, while genes involved in neural development such as *RAX, TAC3* and *NPTX1* were downregulated (Figure 3C).

**Figure 3.**
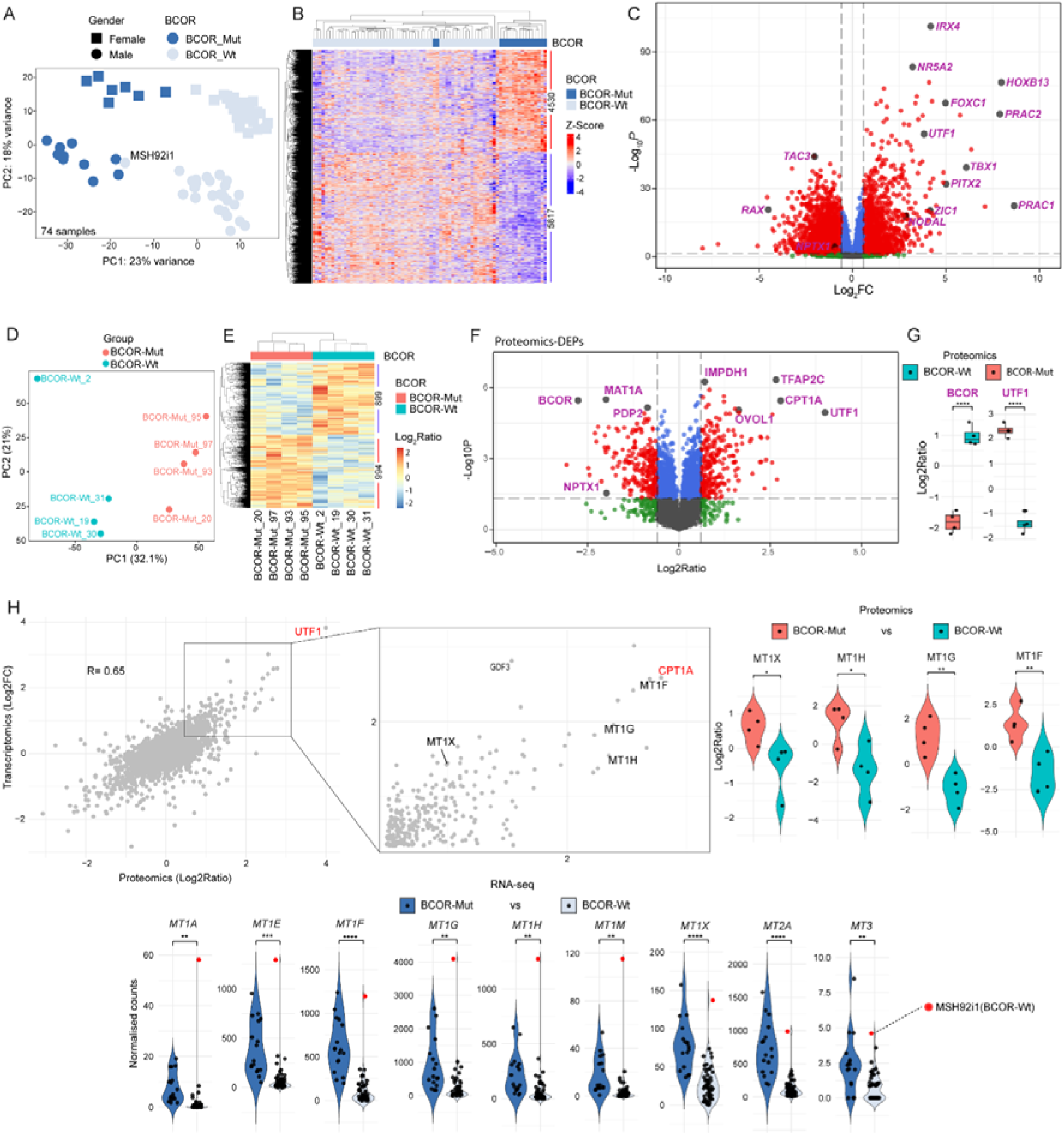
BCOR mutations induce coordinated transcriptomic and proteomic alterations in human iPSCs. **A**, Principal component analysis (PCA) of RNA-seq profiles from B-hiPSCs. Circles and squares denote male and female lines, respectively. **B**, Clustering heatmap of DEGs between BCOR-Mut and BCOR-Wt B-hiPSCs. A total of 4,530 genes were upregulated and 5,817 were downregulated (padj < 0.05). Values represent row-scaled expression (Z-scores). **C**, Volcano plot showing DEGs between BCOR-Mut and BCOR-Wt B-hiPSCs. **D**, PCA of proteomic profiles from four BCOR-Wt and four BCOR-Mut B-hiPSC lines. **E**, Clustering heatmap of differentially abundant proteins (DEPs). A total of 689 proteins were increased and 994 were decreased in BCOR-mutant hiPSCs relative to BCOR-Wt controls (pval < 0.05). **F**, Volcano plot of DEPs between BCOR-mutant and BCOR-Wt. **G**, Relative abundance of BCOR and UTF1 proteins in BCOR-Wt and BCOR-mutant B-hiPSCs. **H**, Correlation between transcriptomic and proteomic fold changes for genes detected in both datasets. **I-K** Normalized RNA-seq counts and relative protein abundance for selected metallothionein genes in BCOR-Wt and BCOR-Mut B-hiPSCs. Individual points represent independent biological replicates. Statistical significance was assessed using a two-sided T-test (*p* < 0.0001****, *p* < 0.001***, *p* < 0.01**, *p* < 0.05*).

Quantitative proteomic analysis of four BCOR-mutant hiPSCs when compared to four BCOR-wildtype lines identified 7,783 proteins with PCA again showing clear separation by genotype (Figure 3D). A total of 1,893 were differentially abundant between groups (pval <0.05), comprising 994 upregulated and 899 downregulated proteins in BCOR-mutant cells (Figure 3E). Pathway enrichment analysis closely mirrored transcriptomic data, with increased representation of oxidative phosphorylation, reactive oxygen species and metabolic pathways in BCOR-mutant cells (Supp Figure 3C, D). Notably, UTF1 protein abundance was significantly increased, whereas BCOR protein levels were markedly reduced in mutant lines.

To assess concordance between molecular strata, we compared RNA-seq fold changes with proteomic abundance changes for overlapping genes. Transcriptomic and proteomic changes were positively correlated (r= 0.65) (Figure 3H), indicating that a substantial proportion of transcriptional changes were translated to the protein level. Among the most consistently upregulated genes across both datasets were *UTF1*, *CPT1A,* and multiple metallothionine family members, including *MT1F*, *MT1G*, *MT1H* and *MT1X*, suggesting enhanced cellular adaptation to metabolic and oxidative stress.

Given the prominent enrichment of mitochondrial metabolic pathways, we next examined genes involved in energy metabolism in greater detail. Both transcriptomic and proteomics analyses demonstrated coordinated upregulation of genes involved in fatty acid β-oxidation, tricarboxylic acid (TCA) cycle, and mitochondrial respiratory chain. In particular, *CPT1A,* the rate-limiting enzyme of fatty acid β-oxidation, was significantly increased in BCOR-mutant hiPSCs. Increased expression was also observed for several Voltage-Dependent Anion Channels (VDACs) such as *VDAC1*, *VDAC2*, *VDAC3,* (Figure 4A, B) which facilitate metabolite exchange across the mitochondrial outer membrane.

**Figure 4.**
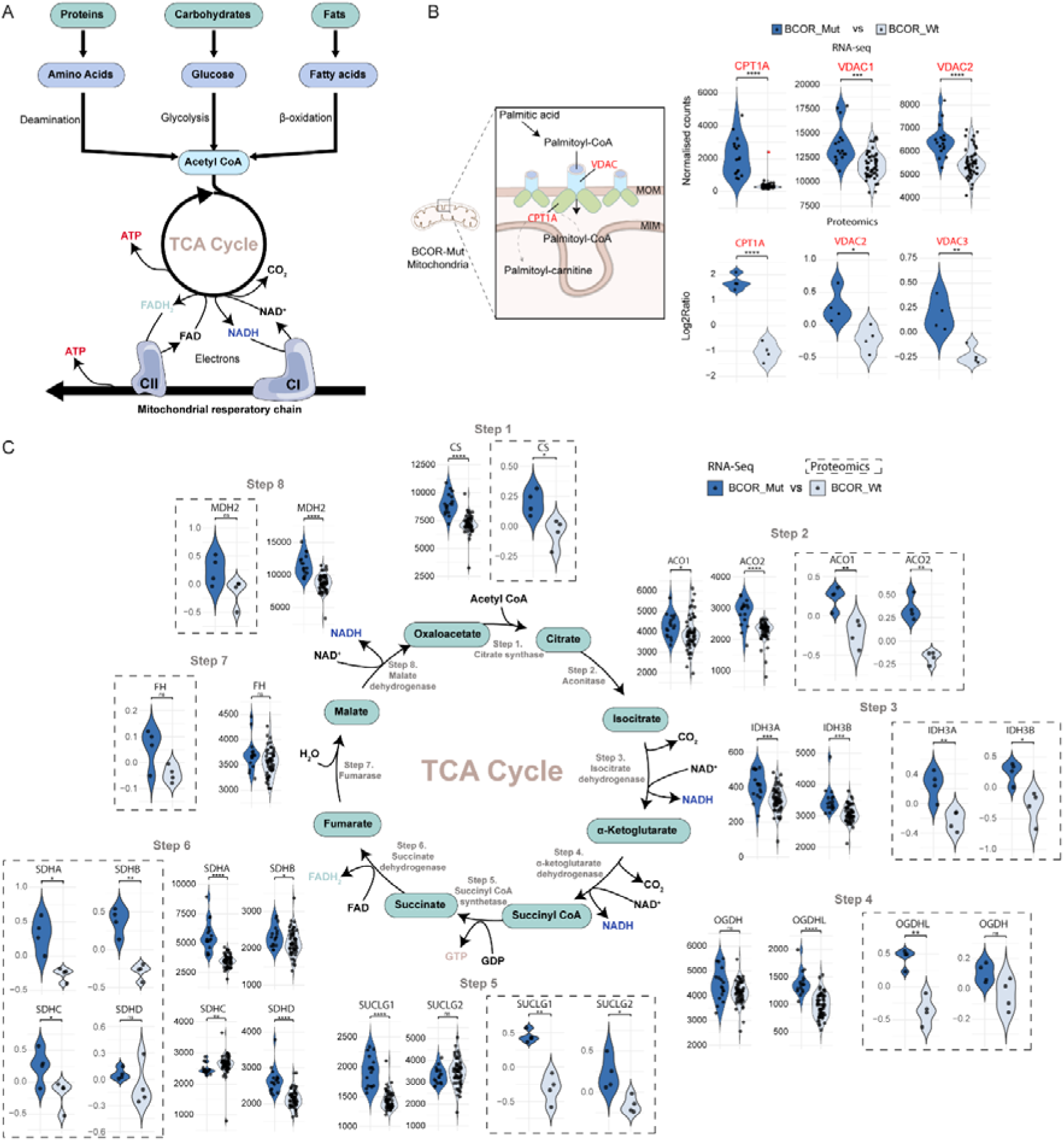
BCOR-mutant hiPSCs exhibit coordinated activation of energy metabolism. **A**, Schematic overview of metabolic pathways altered in BCOR-mutant hiPSCs based on integrated transcriptomic and proteomic analyses. **B**, Expression of key regulators of mitochondrial fatty acid transport and β-oxidation in BCOR-mutant and BCOR-wildtype hiPSCs. **C**, Transcriptomic and proteomic expression changes in genes involved in the TCA cycle. Statistical significance was assessed using a two-sided t-test (*p* < 0.0001****, *p* < 0.001***, *p* < 0.01**, *p* < 0.05*, ns = non-significant).

This metabolic shift extended beyond fatty acid oxidation. Nearly all genes involved in the TCA cycle were significantly upregulated in both omics datasets (Figure 4C). Similarly, multiple components of the mitochondrial electron transport chain were significantly enriched in BCOR-mutant cells (Supp Figure 4B, 4C), indicating increased capacity for oxidative energy production. Multiple genes in the glyconeogenesis/glycolysis were dysregulated (Supp Figure 4A) which has not been reported before(Puigdevall *et al*., 2023; Rouhani *et al*., 2022; Wang *et al*., 2018).

### Multi-omics integration identifies a mitochondrial metabolic reprogramming as a central consequence of BCOR loss

To determine whether the chromatin, transcriptomic, and proteomic alterations observed in BCOR-mutant hiPSCs reflected coordinated regulation across molecular layers, we performed integrative multi-omics analysis using Multi-Omics Factor Analysis 2 (MOFA2)(Argelaguet et al., 2018). This unsupervised approach decomposes variance across datasets into latent factors allowing the identification of shared sources of biological variation (Figure 5A).

**Figure 5.**
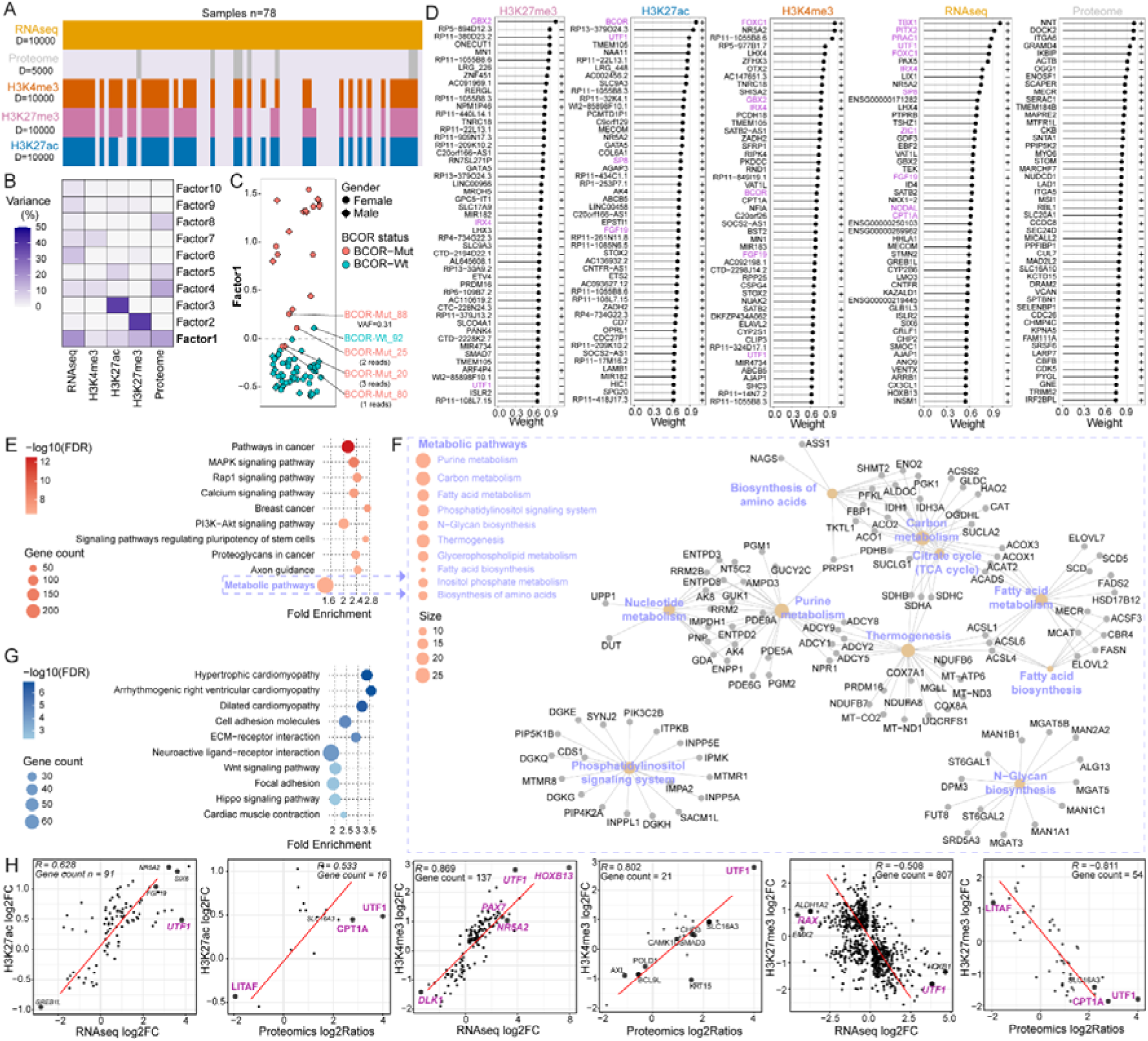
Multi-omic integration identifies a coordinated molecular programme linking chromatin, transcriptional and proteomic alterations. A, Overview of molecular datasets included in the multi-omics factor analysis (MOFA). **B,** Proportion of variance explained by each latent factor across individual molecular layers. **C,** Distribution of samples according to Factor 1. **D,** Top weighted features contributing to Factor 1 across H3K27me3, H3K27ac, H3K4me3, RNA-seq and proteomic datasets. **E,** GO enrichment analysis of positively weighted Factor 1 features. **F,** Network representation of significantly enriched metabolic pathways. **G,** GO enrichment analysis of negatively weighted Factor 1 features. **H,** Correlation of chromatin, transcriptomic and proteomic change identified by supervised differential expression analyses.

Among the ten latent factors identified, only Factor 1 segregated BCOR-mutant from BCOR-Wt hiPSCs (Figure 5B-C, Supp Figure 5A); indicating that BCOR status defines a dominant axis of molecular variation across the integrated datasets. In contrast, the remaining factors captured chromatin-specific and other sources of variance unrelated to genotype (Figure 5B-C, Supplementary Figure 5A). Importantly, Factor 1 received contributions from all molecular layers examined, including H3K27me3, H3K27ac, H3K4me3, RNA-seq and proteomics data. Features with the highest Factor 1 weights included UTF1, FOXC1, IRX4, NR5A2, PITX2, ZIC1, GBX2 and CPT1A (Figure 5D), highlighting regulators of pluripotency, developmental patterning and metabolism that had also emerged from the individual omics analyses.

Functional enrichment of the Factor 1 features revealed strong enrichment of oxidative phosphorylation, fatty acid metabolism, the TCA cycle, carbon metabolism and nucleotide biosynthesis pathways (Figure 5E, 5F) whereas pathways related to cell adhesion, extracellular-matrix organisation and neuroactive signalling were depleted (Figure 5G). These findings indicate that BCOR mutations drive a coordinated regulatory programme centred on mitochondrial energy metabolism that is detectable across chromatin, transcriptomic and proteomic layers. The same metabolic axis emerges in BCOR-mutant AML cell lines(Robert et al., 2026), suggesting that this multi-layered metabolic rewiring is a conserved consequence of BCOR loss across pluripotent and haematopoietic contexts.

To directly assess concordance between molecular layers, we next performed a supervised differential expression analysis and compared the lists of differentially expressed genes (DEGs) and proteins (DEPs) to the closest genes associated with altered -H3K27me3, H3K27ac, H3K4me3-deposition. Consistent with the MOFA2 analysis, genes associated with active chromatin marks (H3K27ac and H3K4me3) showed positive correlations with both transcript and protein abundance, whereas genes associated with repressive marks H3K27me3 showed inverse relationships (Figure 5H, Supp Figure 5D-5J). Shared upregulated features included *UTF1*, *CYGB*, *SLC16A3*, and *CPT1A*, while *CADM4*, *LITAF*, *SARM1*, *ATP1A2*, and *NOVA2* were consistently downregulated across multiple datasets. Together, these analyses demonstrate that BCOR mutations drive a coherent multi-layered regulatory programme linking chromatin remodelling to transcriptional and proteomic states, with metabolomic reprogramming emerging as a central consequence of BCOR loss.

### A qPCR-based assay enables rapid detection of BCOR-mutant hiPSCs

Given the widespread molecular remodelling associated with BCOR mutations, their impact on differentiation potential(Puigdevall *et al*., 2023; Rouhani *et al*., 2022), and the continued emergence of BCOR-mutant clones during routine culture(Rouhani *et al*., 2022), we sought to develop a rapid and cost-effective assay for monitoring BCOR status without requiring genome-wide profiling.

Using our RNA-seq discovery dataset, we defined a fifteen gene biomarker panel comprising three pluripotency markers (*NANOG*, *POU5F1*, *SOX2*), two ncPRC1.1 components (*BCOR* and *KDM2B)* seven consistently upregulated genes (*FOXC1*, *UTF1*, *ZIC1*, *PRAC1*, *PITX2*, *TBX1* and *NODAL*) and three downregulated neural-development genes (*NPTX1*, *RAX* and *TAC3*) (Figure 6A). Principal component analysis using only these fifteen genes retained clear separation between BCOR-mutant and BCOR-wildtype hiPSCs, accounting for 56.8% of the variance along PC1, indicating that this reduced panel captures the major genotype-discriminating signal (Figure 6B).

**Figure 6.**
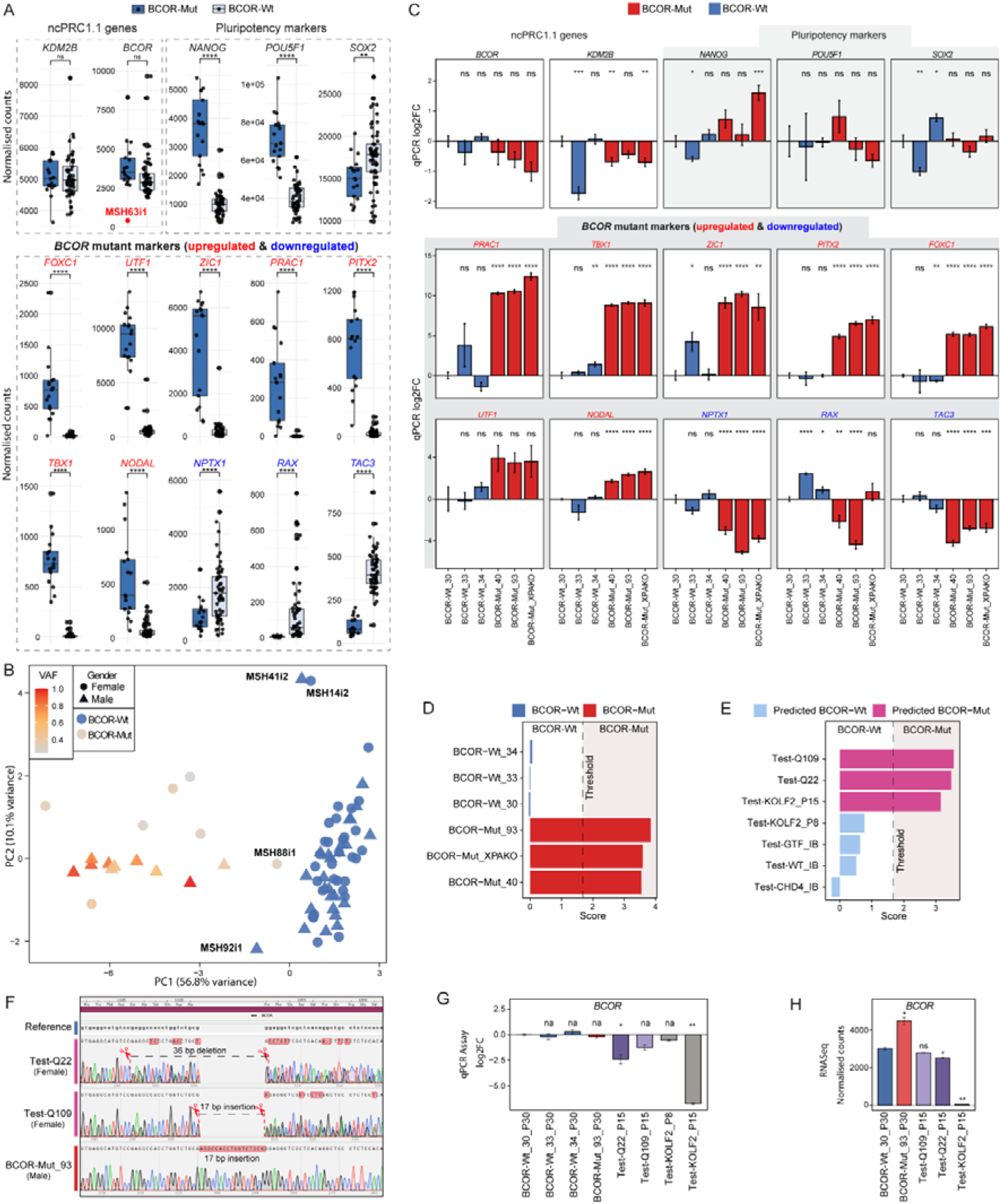
Development and validation of a q-PCR assay for detection of BCOR-mutant hiPSCs. **A**, RNA-seq expression of the 15-gene biomarker panel used for BCOR-mutant detection. Individual points represent independent B-hiPSC lines. Statistical significance was assessed using a two-sided t-test (*p* < 0.0001****, *p* < 0.001***, *p* < 0.01**, *p* < 0.05*, ns = non-significant). **B**, PCA of B-hiPSC lines using expression of the 15-gene biomarker panel. Samples are coloured according to BCOR status and shaded according to BCOR mutant variant allele frequency (VAF). **C**, Validation of the qPCR BCOR-mutant detection assay in independent B-hiPSC lines. Expression values are shown as log_₂_ fold change relative to the BCOR-Wt_30 sample. **D**, BCOR transcriptional scores calculated from the qPCR assay in three reference BCOR-wiltype and three reference BCOR-mutant lines. **E**, Application of the qPCR assay on independent test cohort of hiPSC lines. **F**, Sanger sequencing validation of BCOR mutations identified in qPCR-predicted BCOR-mutant lines. **G**,**H**, BCOR expression in reference and test lines by qPCR (G) and RNA-seq (H). Values are shown relative to BCOR-Wt_30_P30. Statistical significance was assessed using a two-sided t-test (*p* < 0.0001****, *p* < 0.001***, *p* < 0.01**, *p* < 0.05*, ns = non-significant).

Although BCOR mutations are predicted loss-of-function events and result in reduced protein abundance (Figure 3C, 3F-G), *BCOR* transcript levels alone were insufficient to distinguish mutant from wild-type cells. We therefore focused on downstream transcriptional biomarkers that capture the functional consequences of *BCOR* loss. Consistent with this approach, transcriptomic signatures from *KDM2B*-knockout (*KDM2B*-KO) human embryonic stem cells(Wang *et al*., 2018) positively correlated with both our BCOR-mutant RNAseq (R = 0.434, n= 506) (Supp Figure6A top) and proteomic data (R= 0.52, n=124) (Supp Figure6A bottom, 6B) suggesting that much of the observed phenotype reflects disruption of ncPRC1.1 function.

To translate this signature into a practical screening tool, we quantified all fifteen markers using commercially available TaqMan assays (ThermoFisher). Across six fully characterised hiPSC lines, three BCOR-Wt (BCOR-Wt_30, BCOR-Wt_33, BCOR-Wt_34) and three BCOR-mutant (BCOR-Mut_40, BCOR-Mut_93, BCOR-Mut_XPAKO), qPCR-derived expression profiles closely recapitulated the RNA-seq results (Figure 6C). Pluripotency markers remained stable across genotypes, whereas BCOR-mutant signature genes showed the expected upregulation and neural-development markers were consistently reduced (Figure 6C).

We next derived a single per-sample BCOR-Mut score and classification threshold from these reference lines, of 1.67. Assuringly, all BCOR-Mut samples scored above the threshold, whereas all BCOR-Wt samples scored below it (Figure 6D).

To evaluate assay performance in an independent setting, we applied the same scoring framework to hiPSC lines obtained from multiple laboratories (Figure 6E). The Test-Q109 is a Huntington’s disease hiPSC model carrying 109 CAG repeats in the HTT gene and is derived from a female individual, while Q22 is an isogenic corrected derivative generated from Q109(Stöberl et al., 2023). The KOLF2 line, also derived from a female individual, is a widely used reference hiPSC line(Gracia-Diaz et al., 2024; Pantazis et al., 2022; Ryan et al., 2024) (Figure 6 E). Our assay classified Test-GTF_IB, Test-CHD_IB, Test-Wt_IB hiPSCs and early passage KOLF2 (P8) as BCOR-Wt, while Test-Q109, Test-Q22 and Test-KOLF2_P15 were classified as BCOR-mutant (Figure 6E, Supp Figure 6C). These classifications were independently supported by bulk RNA-seq data of the same samples (Supp Figure 6D).

To further validate these results, lines classified as BCOR-mutant were subjected to targeted Sanger sequencing. Consistent with the qPCR results, both Q109 and Q22 harboured exon 7 BCOR mutations (Figure 6F). Notably, the two lines carried distinct indels; where Q109 harboured a 17-nucleotide insertion (AGCCACCTGGTCTGCGA), while Q22 (a derivative of Q109) had a 19-nucleotide deletion (GAGGCCACCTGGTCTGCGG) adjacent to the insertion site in Q109. These observations are consistent with ongoing mutational evolution. That an already BCOR mutant line can undergo further BCOR mutations suggests that the mutability of this region is high in hiPSCs and/or the selective pressure to maintain BCOR dysregulation is substantial.

In contrast, no BCOR mutation was detected in Test-KOLF2_P15 despite its classification as BCOR-mutant. However, both qPCR and RNA-seq revealed markedly reduced BCOR transcript abundance relative to early-passage KOLF2 and other tested lines (Figure 6G,H; Supplementary Figure 6C,D), suggesting that alternative mechanisms, such as structural variation or epigenetic silencing, may disrupt BCOR function in this line. Altogether, these findings establish a rapid, low-cost assay for routine surveillance of BCOR status during hiPSC culture and clonal expansion.

### CRISPR correction of BCOR exon 7 induces incomplete restoration of the BCOR-wild-type molecular state

Having established that BCOR mutations drive extensive molecular remodelling in hiPSCs and can be detected using our qPCR assay, we next asked whether BCOR-mutant lines could be genetically rescued. If correction of the mutant allele restores a BCOR-wildtype molecular state, existing BCOR-mutant cultures could potentially be salvageable for downstream applications. We therefore tested whether precise CRISPR-Cas9 correction of a recurrent exon 7 BCOR mutation was sufficient to restore the BCOR-mutant phenotype.

We selected the male B-hiPSC line BCOR-Mut_93, which carries a recurrent 17-nucleotide insertion in exon 7 that introduces a frameshift and premature stop codon. To selectively revert this allele to wildtype, we designed an allele-specific CRISPR-Cas9 strategy in which the sgRNA spans the insertion junction, a sequence absent from wildtype BCOR alleles (Figure 7A). Homology-directed repair using a single-stranded donor template restores the wildtype exon 7 sequence and eliminates the sgRNA recognition site, thereby preventing repeated cleavage.

**Figure 7.**
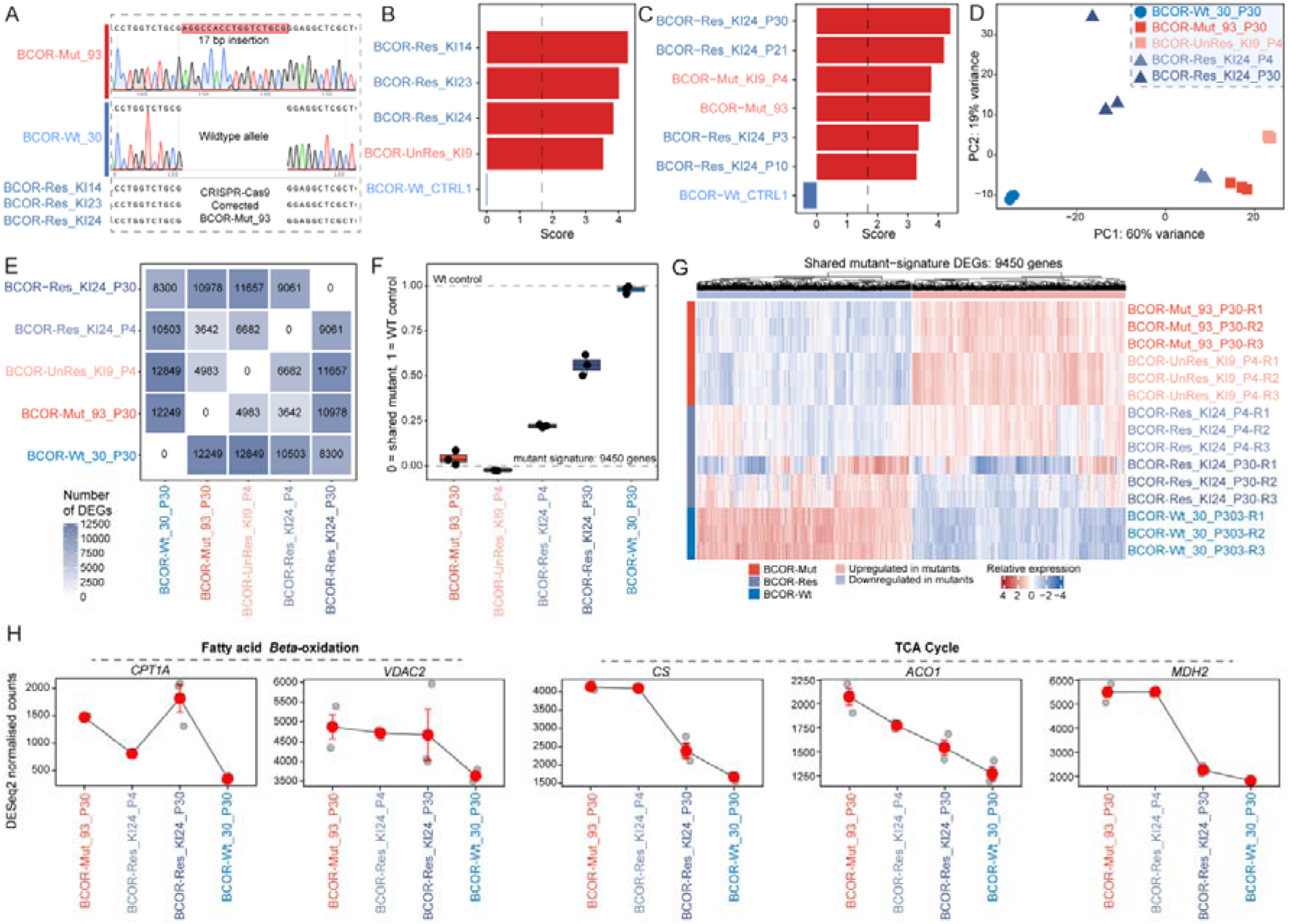
CRISPR correction of BCOR induces incomplete restoration of the BCOR-wildtype transcriptional programme. **A**, Sanger sequencing validation of CRISPR-Cas9 correction of a 17-bp exon 7 insertion in BCOR-Mut_93 hiPSCs. **B**,**C**, BCOR classifier scores generated using the qPCR-based assay in corrected (BCOR-Res), uncorrected (BCOR-UnRes), mutant (BCOR-Mut) and control (BCOR-Wt) lines. Dashed lines indicate the classification threshold. **D**, PCA of RNAseq profiles mutant, rescued, uncorrected and wildtype BCOR clones. **E**, Pairwise differential expression analysis between BCOR-Wt, BCOR-Mut, BCOR-UnRes and BCOR-Res samples. Numbers indicate the total number of DEGs identified in each comparison. **F**, Quantification of the 9,450-gene BCOR-mutant transcriptional signature. Values are scaled from 0 (BCOR-mutant state) to 1 (BCOR-wildtype state). **G**, Heatmap showing expression of the shared 9,450-gene BCOR-mutant transcriptional signature across BCOR-Wt, BCOR-Mut, BCOR-UnRes and BCOR-Res samples. **H**, RNA-seq expression of representative genes involved in fatty-acid β-oxidation (*CPT1A*, *VDAC2*) and the tricarboxylic acid cycle (*CS*, *ACO1*, *MDH2*) in BCOR-Mut_93_P30, BCOR-Res_KI24_P4, BCOR-Res_KI24_P30 and BCOR-Wt_30_P30. Points represent individual biological replicates and red symbols indicate group means.

Sanger sequencing confirmed precise correction of the mutant allele in three independent subclones (BCOR-Res_KI14, BCOR-Res_KI23 and BCOR-Res_KI24)(Figure 7A). Surprisingly, however, all corrected clones remained above the predefined BCOR-mutant threshold when assessed using the qPCR classifier. All scored similarly to the parental BCOR-Mut_93 and a non-rescued edited clone BCOR-UnRes_KI9 (Figure 7B). Longitudinal monitoring of BCOR-Res_KI24 over multiple passages likewise showed persistent classification as BCOR-mutant state despite successful genetic correction (Figure 7C).

To determine whether correction induced broader transcriptional changes, we performed RNA-seq on BCOR-Mut_93, BCOR-UnRes_KI9, BCOR-Res_KI24 at early passage (P4) and late passage (P30), and a BCOR-Wt_30 late passage control. Principal component analysis showed that BCOR-UnRes_KI9 clustered closely with BCOR-Mut_93, whereas BCOR-Res_KI24 occupied an intermediate position between mutant and wildtype states (Figure 7D). Notably, the P30 corrected sample shifted further towards the wildtype cluster than the P4 sample, suggesting progressive but incomplete restoration of the transcriptional state.

Differential expression analysis supported this interpretation (Figure 7E). Whereas BCOR-Mut_93 and BCOR-UnRes_KI9 differed substantially from BCOR-wildtype cells, BCOR-Res_KI24 showed a gradual reduction in transcriptional divergence from the wildtype state between P4 and P30 (Figure 7E). To quantify this transition, we defined a shared BCOR-mutant transcriptional signature and projected each sample onto a mutant-to-wildtype continuum. Consistent with the PCA results, corrected cells shifted away from the mutant state following repair, with a greater shift observed at P30 than at P4, but neither sample reached the wildtype endpoint (Figure 7F).

Heatmap analysis of the shared BCOR-mutant signature further demonstrated partial normalisation of both mutant-upregulated and mutant-downregulated genes following correction (Figure 7G,H). However, substantial residual differences persisted relative to BCOR-wild-type cells, even after extended culture.

Together, these findings indicate that precise correction of a recurrent BCOR exon 7 mutation is insufficient to fully restore the wildtype molecular state. Although rescued cells progressively move towards a BCOR-wild-type transcriptional profile, residual mutant-associated programmes remain detectable, suggesting that prevention and early detection of BCOR-mutant clones may be more effective than attempting genetic rescue after their emergence.

## Discussion

Human induced pluripotent stem cells have become indispensable tools for disease modelling, drug discovery, and regenerative medicine. Their utility, however, depends on models maintaining their molecular integrity during prolonged culture and clonal expansion. In this study, we show that recurrent *BCOR* mutations represent a distinctive form of culture adaptation in hiPSCs, characterised by extensive chromatin, transcriptional and proteomic remodelling, activation of metabolic pathways associated with cellular fitness, and persistence of mutant-associated molecular states even after precise genetic correction.

Our analysis confirms and extends previous reports that *BCOR* mutations are among the most recurrently selected genetic alterations in hiPSCs(Rouhani *et al*., 2022)^27,28^. Unlike BCOR mutations reported in AML and other cancers, hiPSC mutations were strongly enriched for truncating indels within exon 7, indicating that the selective pressures operating in pluripotent stem cells differ from those acting during tumorigenesis. The observation of distinct sequential mutational events affecting the same exon 7 region in the Q109/Q22 isogenic pair further supports the existence of a mutational hotspot in hiPSCs. The co-localisation of recurrent mutations with predicted G-quadruplex sequences and a pluripotent stem cell-specific DNase I hypersensitive site provides a plausible mechanistic explanation for this phenomenon, although direct experimental validation will be required to determine whether local chromatin architecture contributes to mutagenesis at this locus.

Beyond defining the mutational landscape, our study provides insight into the molecular consequences of BCOR loss in hiPSCs. Integration of chromatin profiling, transcriptomics, proteomics and multi-omics factor analysis consistently identified BCOR status as a dominant source of molecular variation. While developmental and differentiation pathways were prominently affected, one of the most striking findings was the coordinated activation of mitochondrial energy metabolism, including fatty acid β-oxidation, the tricarboxylic acid cycle and oxidative phosphorylation. Similar metabolic programmes have been reported following BCOR loss in haematopoietic malignancies^56^, suggesting that metabolic rewiring may represent a conserved consequence of BCOR disruption across cellular contexts. Importantly, the enhanced survival of BCOR-mutant nucleotide excision repair-deficient hiPSCs following UV-induced DNA damage provides functional support for the hypothesis that these molecular changes confer a selective advantage under stress conditions. Together, these observations suggest that BCOR mutations are not merely tolerated during culture but actively promote cellular states that favour survival and expansion.

A key translational outcome of this work is the development of a simple qPCR-based assay capable of detecting BCOR-mutant transcriptional states across independent hiPSC models. Current quality-control pipelines often rely on periodic karyotyping or targeted genotyping, approaches that may fail to identify emerging mutant subclones before they become dominant within a culture. By capturing the downstream transcriptional consequences of BCOR dysfunction rather than mutation status alone, our assay provides a practical and inexpensive approach for routine surveillance during long-term culture, clonal expansion and genome editing workflows. The identification of a BCOR-dysregulated KOLF2 subclone lacking detectable coding mutations further illustrates the value of functional readouts that extend beyond sequence-based screening alone.

Finally, our CRISPR rescue experiments revealed that precise correction of a recurrent BCOR exon 7 mutation induces only partial restoration of the wildtype molecular state. Although corrected cells progressively shifted towards a BCOR-wildtype transcriptional profile over time, residual mutant-associated signatures remained detectable after extended culture. These findings are consistent with emerging evidence that transient disruption of Polycomb-mediated regulation can establish self-reinforcing epigenetic states that persist after the initiating lesion has been removed(Parreno et al., 2024). From a practical perspective, this observation has important implications for stem cell quality control. If restoration of the wildtype state requires prolonged culture, the opportunity for additional adaptive mutations to arise may increase, potentially limiting the utility of post hoc genetic correction as a remediation strategy.

Taken together, our findings establish BCOR mutations as a major source of molecular divergence in hiPSC cultures and identify metabolic reprogramming as a central consequence of BCOR loss. More broadly, they highlight the need for routine molecular surveillance of hiPSC lines and suggest that prevention and early detection of adaptive variants may be more effective than attempting to reverse their consequences once established.

## Limitations of the study

Several limitations should be acknowledged. First, although the molecular consequences of BCOR loss were characterised across multiple omics layers, our conclusions regarding metabolic rewiring are based primarily on chromatin, transcriptomic and proteomic data and were not complemented by direct measurements of metabolic flux, mitochondrial respiration or ATP production. Second, while the recurrent enrichment of mutations within exon 7, particularly at predicted G-quadruplex sequences, is consistent with the existence of a mutational hotspot, the underlying mechanisms driving recurrent mutagenesis at this locus remain to be experimentally validated. Finally, although our CRISPR rescue experiments demonstrate incomplete restoration of the wildtype molecular state following BCOR correction, these studies were performed on a single recurrent exon 7 mutation, and it remains possible that other BCOR mutations, genetic backgrounds or culture conditions may exhibit different rescue dynamics.

## Conclusion

Two decades after the successful generation of the first iPSC line, the technology has matured to the point where its limitations matter as much as its capabilities. BCOR mutations exemplify a class of culture-acquired alterations that are individually small, easily missed by conventional surveillance, irreversible by genetic correction once established, and consequential for the very neural and developmental applications hiPSCs are most often used to model. The framework and assay presented here address one specific case, but the broader lesson generalises, ensuring the integrity of hiPSC models requires sustained, signature-level vigilance throughout the life of a line, not a one-time certification at its origin. As the field continues to expand into the clinic, developing tools and standards that improve the safety and quality control of hiPSCs represents, in our view, one of the most important contributions basic research can make to the translational promise of induced pluripotency.

## Acknowledgments

We would like to thank the Sequencing Operations team at the Wellcome Sanger Institute (Hinxton, UK), and the National Xeroderma Pigmentosum Service, St John’s Institute of Dermatology, Guy’s and St Thomas’ Foundation Trust (London, UK). We extend our gratitude to our funders: Sir Jeffery Cheah Foundation (GCCK); Cancer Research UK Advanced Clinician Scientist Award (C60100/A23916) (SNZ); Dr. Josef Steiner Cancer Research Award 2019; Basser Gray Prime Award 2020 (SNZ); CRUK Pioneer Award (C60100/A23433) (SNZ); CRUK Grand Challenge Award (C60100/A25274) (SNZ); CRUK Early Detection Project Award (C60100/A27815) (SNZ); National Institute for Health and Care Research (NIHR) Professorship (NIHR301607) (SNZ); NIHR Cambridge Biomedical Research Centre (BRC-1215-20014, NIHR203312) (SNZ). The views expressed are those of the authors and not necessarily those of the NIHR or the Department of Health and Social Care.

## Supplementary

Supplementary tables are provided as an Excel spreadsheet.

## METHOD DETAILS

### Establishing human induced pluripotent stem cells

Human induced pluripotent stem cell (hiPSC) models were generated through the Cellular Generation and Phenotyping facility at the Wellcome Sanger Institute. Donor peripheral blood mononuclear cells (PBMCs) were isolated from total blood, erythroblasts were expanded, and cells were reprogrammed using the four Yamanaka factors OCT3/4, SOX2, KLF4 and c-MYC. Reprogramming was performed using the non-integrating CytoTune Sendai Reprogramming Kit (Invitrogen), according to the manufacturer’s instructions.

All hiPSC lines used in this study underwent standard quality-control assessment, including chromosomal copy-number profiling, immunofluorescence staining for pluripotency markers and expression-based assessment of pluripotency. Only lines showing typical undifferentiated hiPSC colony morphology and meeting routine quality-control criteria were included in the study.

### hiPSC maintenance

All hiPSC lines at the Nik-Zainal lab were maintained under feeder-free conditions on Vitronectin FX-coated plates (STEMCELL Technologies) in Essential 8 medium (Gibco). Medium was replaced daily and cultures were routinely inspected to ensure a healthy undifferentiated morphology without signs of spontaneous differentiation. For routine propagation, hiPSCs were washed with DPBS (Gibco), treated with 0.5 mM UltraPure EDTA (Invitrogen), and passaged as small cell clumps onto freshly coated plates. Whenever single-cell dissociation was required (e.g. CRISPR-Cas9 editing or colony-formation assays), cells were detached using TrypLE Express Enzyme (Gibco) and replated in Essential 8 medium supplemented with RevitaCell Supplement (Gibco).

### Culture conditions for independent validation hiPSC lines

Independent validation hiPSC lines obtained from collaborators were maintained in different culture conditions. The Q109 and Q22 lines were cultured in StemMACS iPS-Brew medium (Miltenyi Biotec) supplemented with 1/100 Penicillin-Streptomycin, with daily medium changes. Cells were maintained on Bio-Techne Cultrex PathClear Basement Membrane Extract (BME), diluted 1/100 in calcium- and magnesium-free DPBS and coated overnight at 4°C. Coated plates were incubated at 37°C for 30 min before use. Cells were passaged when approximately 70% confluent using 0.5 mM EDTA in calcium- and magnesium-free DPBS for 8 min at 37°C. Cells were cryopreserved in freezing medium containing 90% KnockOut Serum Replacement and 10% DMSO. KOLF2 lines were cultured in StemMACS iPS-Brew medium (Miltenyi Biotec) supplemented with 1/100 Penicillin-Streptomycin, with daily medium changes. Cells were maintained on Biolamina Laminin 521, diluted in calcium- and magnesium-containing DPBS and coated overnight at 4°C. Coated plates were incubated at 37°C for 30 min before use. Cells were passaged when approximately 70% confluent using 0.5 mM EDTA in calcium- and magnesium-free DPBS for 8 min at 37°C. Cells were cryopreserved in freezing medium containing 90% KnockOut Serum Replacement and 10% DMSO. Test-WT_IB, Test-GTF_IB and Test-CHD4_IB were maintained on vitronectin-coated culture vessels in Essential 8 (E8) medium and cultured at 37°C with 5% CO₂. Medium was refreshed daily, and cells were passaged every 4–5 days at 60–80% confluency using either 0.5 mM EDTA for colony-based passaging or Accumax for single-cell dissociation when required. Single-cell cultures were supplemented with 10 μM Y-27632 ROCK inhibitor or 1× RevitaCell for the first 24 h following passaging.

### Compilation and annotation of BCOR mutations

Reported BCOR mutations in hiPSCs were compiled from previously published hiPSC genomic datasets and compared with BCOR mutations reported in acute myeloid leukaemia (AML) and non-AML cancer cohorts. Cancer mutation data were obtained from cBioPortal. BCOR variants from cBioPortal were curated according to cancer type annotation and grouped into AML and non-AML cancer cohorts for comparison with the hiPSC mutation dataset. Variants were assigned to BCOR exons using the canonical BCOR transcript annotation. Variant consequences were classified as frameshift insertion/deletion, stop-gained, missense or other coding consequence. The number and proportion of variants in each exon were calculated separately for hiPSC, AML and non-AML cancer cohorts. Exon-level enrichment was assessed using Fisher’s exact tests by comparing the number of mutations in each exon against mutations in all other BCOR exons within the relevant comparator cohorts.

### BCOR sequence-context and G-quadruplex analysis

BCOR exon sequences and flanking genomic regions were extracted from the human reference genome. The local sequence context of each reported BCOR mutation was inspected to determine whether variants occurred at, or immediately adjacent to, G/C nucleotides. Recurrent inserted or deleted motifs in exon 7 were aligned to the BCOR reference sequence to define the recurrent mutation hotspot region.

Putative G-quadruplex-forming sequences were predicted using QGRS Mapper. Predicted motifs with a G-score greater than 30 were retained for downstream analysis. Predicted G-quadruplex motifs were intersected with BCOR exon coordinates and plotted relative to mutation positions. DNase I hypersensitive site annotations from pluripotent stem cell datasets available on UCSC (deposited by the Duke University) were used to identify open chromatin regions overlapping predicted G-quadruplex motifs within BCOR.

### ChIPmentation and ChIP-seq library preparation

Chromatin profiling was performed using a ChIPmentation-based protocol for formaldehyde-crosslinked hiPSCs. The histone marks profiled were H3K4me3, H3K27ac and H3K27me3. Crosslinked hiPSC pellets were lysed, sonicated, immunoprecipitated using histone-mark-specific antibodies and processed into Illumina-compatible sequencing libraries using a tagmentation-based workflow.

All lysis and chromatin-handling steps were performed under chilled conditions to preserve protein-chromatin interactions. Crosslinked pellets were thawed on ice and sequentially lysed using Diagenode iL1 and iL2 buffers. Nuclei were then incubated in iS1 buffer supplemented with protease inhibitor. Chromatin was transferred to pre-chilled Biorupter microtubes and sheared using a Picorupter for five cycles of 30 s on and 30 s off. The lysate was centrifuged at 16,000 g for 10 min at 4°C and the clarified chromatin-containing supernatant was retained. A small aliquot was removed for shearing assessment where required, and the remaining material was frozen on dry ice and stored at −80°C.

For immunoprecipitation, sheared chromatin was combined with 5% BSA, protease inhibitor, water, 5x iC1 buffer and the appropriate antibody. The antibodies used were Diagenode H3K4me3 ChIP-seq antibody Classic (C15200152), Diagenode H3K27ac ChIP-seq antibody Classic (C15410174) and Diagenode H3K27me3 ChIP-seq antibody Classic (C15410069). For each immunoprecipitation reaction, 75 µL sheared chromatin was used with 1 µg antibody for H3K4me3 and H3K27ac, and 2 µg antibody for H3K27me3. DiaMag protein A-coated beads were washed in chilled iC1 buffer and added to each reaction. Immunoprecipitation was performed overnight at 4°C with shaking.

Following immunoprecipitation, beads were washed sequentially with iW1, iW2, iW3, 10 mM Tris pH 8 and iW4 buffers. On-bead tagmentation was performed using freshly prepared tagmentation buffer and Tn5 enzyme for 10 min at 37°C with shaking. Following tagmentation, beads were washed again and chromatin was eluted in iE1 buffer. Samples were reverse-crosslinked at 65°C for at least 4 h or overnight.

Input samples were processed manually in parallel because they did not undergo antibody-mediated protein A bead pull-down. For each input sample, 2 µL of sheared chromatin was used as template for tagmentation, followed by reverse crosslinking with NaCl, RNase A, proteinase K and iE1 buffer.

Libraries were purified using SPRI bead clean-up, amplified using KAPA HiFi ready mix and indexed with i5 and i7 primers. PCR cycling consisted of 72°C for 5 min, 98°C for 30 s, amplification cycles of 98°C for 10 s, 63°C for 30 s and 72°C for 1 min, followed by a final extension at 72°C for 3 min. Post-PCR libraries were size-selected using double-SPRI clean-up and prepared for Illumina sequencing.

### ChIP-seq analysis

The 150 bp paired-end ChIP-seq reads were mapped to the hg19 human genome assembly using Bowtie 2 v2.3.2. BigWig tracks were generated using the bamCoverage program from the deepTools software suite v2.5.3. Reads were extended to 300 bp and tracks were normalised using counts per million mapped reads (CPM).

Peak calling was performed using MACS2 v2.1.1.20160309. For most ChIP-seq datasets, default MACS2 settings were used. Artefactual peaks overlapping blacklisted genomic regions defined by the Human ENCODE project were removed using bedtools intersect from BEDTools v2.26.0.

Promoter regions were defined as +/-2 kb around annotated transcription start sites. For downstream analyses of promoter and regulatory regions, the top 10,000 peaks for each histone mark were retained. H3K4me3 peaks were used to represent promoter-associated active chromatin, H3K27ac peaks were used to represent active regulatory chromatin, and H3K27me3 peaks were used to represent Polycomb-associated repressive chromatin. Peaks were mapped to the nearest gene using bedtools closest.

For ChIP-seq heatmaps and profile plots, signal matrices were generated using computeMatrix from deepTools v2.5.3 and visualised using plotHeatmap. Regions spanning −2 kb to +2 kb around the centre of each peak were used. These regions were divided into 10 bp or 50 bp bins, and each bin was assigned the mean BigWig signal score across that interval. Heatmaps and average profile plots were generated separately for H3K4me3, H3K27ac and H3K27me3.

For differential chromatin analyses, peak-level count matrices were generated separately for each histone mark and compared between BCOR-mutant and BCOR-wildtype hiPSCs. Differentially enriched or depleted regions were annotated to the nearest gene and used for downstream pathway enrichment and integration with RNA-seq and proteomic datasets.

### RNA extraction and RNA sequencing

Total RNA was extracted using the PureLink RNA Mini Kit (ThermoFisher) according to the manufacturer’s recommendations. RNA quality was assessed before library preparation. cDNA libraries were prepared from high-quality RNA and sequenced using Illumina NovaSeq 6000 technology. For each sample, more than 20 million 150 bp paired-end read pairs were generated.

For the CRISPR rescue experiment, RNA-seq was performed on the parental BCOR-mutant line BCOR-Mut_93, the non-rescued edited clone BCOR-UnRes_KI9, the rescued clone BCOR-Res_KI24 at early passage and at late passage, and a BCOR-wildtype control line BCOR-Wt_30.

### RNA-seq analysis

Splice-aware STAR v2.5.0a was used to map RNA-seq reads to the reference genome. A genome index was generated for the human decoy reference genome hs37d5.fa.gz. FASTQ files were mapped using splice-junction information from the GENCODE v19 GTF annotation file. Reads overlapping gene features were counted using featureCounts v2.0.1.

Raw count matrices were imported into R v4.0.4 and analysed using DESeq2. Lowly expressed genes were removed before differential expression analysis. Differential gene expression was assessed using DESeq2 Wald tests. Genes with adjusted P value < 0.05 were considered differentially expressed unless otherwise stated. Normalised counts and variance-stabilised expression values were used for principal component analysis, heatmaps, sample clustering, correlation analyses and downstream multi-omics integration.

For the main BCOR-mutant versus BCOR-wildtype comparison, BCOR status was modelled as the primary variable. For the CRISPR rescue analysis, normalised expression values were used to compare parental mutant, non-rescued edited, rescued and wildtype states. A shared BCOR-mutant transcriptional signature was used to project rescued samples along a mutant-to-wildtype continuum.

### Proteomics

Lysis buffer stock solution was made using 1 M triethylammonium bicarbonate (TEAB), 5% sodium deoxycholate, 10% isopropanol and 5 M NaCl in water. A lysis buffer working solution was made by adding 2 mL of the lysis buffer stock solution and 20 µL Halt Protease and Phosphatase Inhibitor Single-Use Cocktail (ThermoFisher) to an Eppendorf tube. A total of 150 µL of lysis buffer working solution was added to each sample containing a PBS-washed cell pellet of approximately 3 x 10^6 cells. Samples were heated at 90°C for 5 min and sonicated for a further 5 s. Protein concentration was measured using a Bradford assay with Pierce Coomassie Plus reagent according to the manufacturer’s instructions.

Aliquots containing 30 µg total protein in 20 µL were reduced by adding 2 µL of 50 mM tris-2-carboxyethyl phosphine (TCEP) and incubated in a heat block at 60°C for 1 h. Samples were spun down and alkylated with 1 µL of freshly prepared 200 mM iodoacetamide. The solutions were vortexed, spun down and incubated at room temperature for 30 min in the dark. Trypsin digestion was performed by adding 11 µL of 100 mM TEAB and 6 µL trypsin solution at 500 ng/µL in 0.1% formic acid. Samples were vortexed, spun down and left overnight at room temperature for trypsin digestion. The protein-to-trypsin ratio was approximately 33:1.

A total of 60 µL of 100 mM TEAB was added to each sample to bring the volume to 100 µL. Samples were labelled with TMTpro 16-plex or TMT 11-plex reagent vials according to the manufacturer’s instructions. A total of 41 µL TMT reagent was added to each sample and incubated for 1 h at room temperature. Hydroxylamine, 8 µL of 5%, was added to quench the reaction and incubated for 15 min. All labelled samples were combined in equal amounts into one tube. A total of 20 µL formic acid was added and samples were centrifuged for 5 min at 10,000 rpm to precipitate sodium deoxycholate. The remaining supernatant was transferred into a clean tube and dried using a centrifugal vacuum concentrator.

Pooled control samples were generated by combining 50 µg total protein from control samples to make a stock pooled control. Thirty micrograms of protein from the stock pooled control were added to separate tubes and made up to 20 µL with lysis buffer before reduction, alkylation, digestion and TMT labelling as described above.

### High-pH reversed-phase peptide fractionation and LC-MS analysis

Peptides were fractionated offline by high-pH reversed-phase chromatography using an XBridge C18 column, 2.1 x 150 mm, 3.5 µm, Waters, on a Dionex UltiMate 3000 HPLC system. Mobile phase A was 0.1% ammonium hydroxide and mobile phase B was 100% acetonitrile with 0.1% ammonium hydroxide. TMT-labelled peptides were fractionated at 0.2 mL/min using the following gradient: 5 min isocratic at 5% B, 35 min gradient to 35% B, 5 min gradient to 80% B, 5 min isocratic at 80% B, followed by re-equilibration to 5% B. Fractions were collected every 42 s, pooled into 28 fractions and vacuum dried.

LC-MS analysis was performed on a Dionex UltiMate 3000 UHPLC system coupled to an Orbitrap Lumos mass spectrometer (Thermo Scientific). Each peptide fraction was reconstituted in 40 µL 0.1% formic acid, and 10 µL was loaded onto an Acclaim PepMap 100 C18 trapping column, 100 µm x 2 cm, 5 µm, at a flow rate of 10 µL/min. Samples were analysed using an EASY-Spray C18 capillary column, 75 µm x 50 cm, 2 µm, at 50°C. Mobile phase A was 0.1% formic acid and mobile phase B was 80% acetonitrile with 0.1% formic acid.

The elution method consisted of a 90 min gradient from 5% to 38% B, followed by 10 min up to 95% B, 5 min isocratic at 95% B, re-equilibration to 5% B in 5 min and 10 min isocratic at 5% B, at a flow rate of 300 nL/min. Precursor ions were selected in the range of 375-1,500 m/z with a mass resolution of 120,000, AGC target of 4 x 10^5 and maximum injection time of 50 ms, using top-speed mode in 3 s. Precursor ions were isolated for CID fragmentation with a quadrupole isolation width of 0.7 Th. Collision energy was 35%, with AGC target of 1 x 10^4 and maximum injection time of 50 ms.

Quantification was obtained at the MS3 level with HCD fragmentation of the top five most abundant CID fragments isolated with synchronous precursor selection. The quadrupole isolation width was 0.7 Th, collision energy was 55%, and AGC target was 1 x 10^5 with a maximum injection time of 105 ms. HCD MS3 spectra were acquired for the mass range 100-500 m/z with 50,000 resolution. Targeted precursors were dynamically excluded for 45 s with 7 ppm mass tolerance.

### Database search and protein quantification

Mass spectra were analysed using Proteome Discoverer 2.4 (Thermo Scientific) with the SequestHT search engine for protein identification and quantification. Precursor and fragment ion mass tolerances were 20 ppm and 0.5 Da, respectively. Spectra were searched for fully tryptic peptides with a maximum of two missed cleavages. Static modifications were designated as TMTpro or TMT 6-plex at peptide N-termini and lysine residues, and carbamidomethylation of cysteine. Dynamic modifications were designated as oxidation of methionine and deamidation of asparagine or glutamine.

Peptide confidence was estimated using the Percolator node and peptides were filtered at q value < 0.01 based on a target-decoy database search. All spectra were searched against reviewed UniProt Homo sapiens protein entries. The reporter ion quantifier node included a TMTpro or TMT 11-plex quantification method with an integration window tolerance of 15 ppm at the MS3 level. Only unique peptides were used for quantification, using protein groups for peptide uniqueness. Only peptides with average reporter signal-to-noise ratio > 3 were used for protein quantification.

### Proteomic data analysis using Perseus

Statistical analysis was performed using Perseus software v2.0.7.0. Samples from each TMT run were normalised to the column sum in Proteome Discoverer, and log2 ratios were computed relative to the average of the pooled control samples for the corresponding set. Tukey’s biweight was subtracted for each run to account for batch effects. Data from all TMT runs were then combined, and average values for all control samples combined were subtracted.

For cases where technical replicates were present, an average value was calculated. Proteins with one or zero valid values were filtered out and samples were normalised by row Z score. Student’s t test with a significance threshold of P < 0.05 was applied to compare two groups. Proteins with a significant difference in abundance between groups were extracted using a threshold of −log10 P value > 1.3 and a Student’s t test difference of > 0.5 or < −0.5, corresponding to approximately 1.4-fold change.

Functional classification and pathway analysis were performed within Perseus using GO cellular component, GO biological process, KEGG and GSEA annotations. Fisher’s exact test with Benjamini-Hochberg FDR threshold of 0.02 was applied to determine enriched pathways. Heatmaps showing proteomics results were generated in R using normalised Z-score-scaled data.

### Cross-omics integration and MOFA2 analysis

Multi-omics integration was performed using Multi-Omics Factor Analysis 2 (MOFA2)(Argelaguet *et al*., 2018). Separate input matrices were generated for H3K27me3, H3K27ac, H3K4me3, RNA-seq and proteomics. Features were filtered to retain informative genes or genomic regions and scaled before model training. The integrated model was trained to infer ten latent factors. The variance explained by each factor was calculated across omics layers, and sample factor values were plotted according to BCOR status.

Factor 1 was identified as the major BCOR-associated axis of variation because it separated BCOR-mutant from BCOR-wildtype hiPSCs and received contributions from all molecular layers. High-weight features from Factor 1 were extracted and used for downstream enrichment analysis.

To assess concordance between molecular layers, RNA-seq log2 fold changes were compared with proteomic abundance changes for genes detected in both datasets. Genes nearest to differentially enriched H3K27ac, H3K4me3 and H3K27me3 peaks were compared with RNA-seq and proteomic changes. Pearson correlation coefficients were calculated to quantify concordance between molecular layers. For active marks, H3K27ac and H3K4me3, positive relationships with transcript and protein abundance were assessed. For the repressive mark H3K27me3, inverse relationships with transcript and protein abundance were assessed.

### Pathway enrichment analysis

Pathway enrichment analysis was performed on gene lists derived from differentially expressed genes, differentially abundant proteins, genes nearest to altered ChIP-seq peaks and high-weight MOFA2 features. Upregulated and downregulated feature sets were analysed separately. KEGG, Gene Ontology and related functional annotation datasets were used to identify enriched biological processes and pathways. Enrichment analyses were performed using ShinyGo 0.85.1 and/or R-based enrichment workflows.

### TaqMan qPCR assay and BCOR-mutant score

A 15-gene TaqMan qPCR panel was designed from the RNA-seq discovery dataset to capture the transcriptional consequences of BCOR loss. The panel included three pluripotency markers, NANOG, POU5F1 and SOX2; two ncPRC1.1-related genes, BCOR and KDM2B; seven genes consistently upregulated in BCOR-mutant hiPSCs, FOXC1, UTF1, ZIC1, PRAC1, PITX2, TBX1 and NODAL; and three genes downregulated in BCOR-mutant hiPSCs, NPTX1, RAX and TAC3. GUSB was used as the endogenous control.

Total RNA was extracted from hiPSCs using the Direct-zol RNA Miniprep Plus kit (Zymo Research) or an equivalent column-based RNA extraction method. cDNA was generated using SuperScript IV VILO reverse transcriptase. Where sufficient RNA was available, 1.5 µg RNA was reverse-transcribed. Gene expression was quantified using commercially available TaqMan assays (ThermoFisher). Ct values were inspected for amplification quality, and technical replicates were averaged after exclusion of failed reactions. Delta Ct values were calculated as Ct_target - Ct_GUSB.

For each gene, DeltaDelta Ct values were calculated relative to the relevant reference condition. Relative expression was represented as -DeltaDelta Ct or log2 fold change. For classifier development, expression values from three BCOR-wildtype reference lines, BCOR-Wt_30, BCOR-Wt_33 and BCOR-Wt_34, and three BCOR-mutant reference lines, BCOR-Mut_40, BCOR-Mut_93 and BCOR-Mut_XPAKO, were used to derive a single per-sample BCOR-Mut score.

Genes expected to increase in BCOR-mutant cells contributed positively to the score, whereas genes expected to decrease in BCOR-mutant cells were direction-adjusted before averaging. A fixed classification threshold of 1.67 was derived from the reference lines. Samples with a score above this threshold were classified as BCOR-mutant-like, whereas samples below the threshold were classified as BCOR-wildtype-like. The same threshold was applied unchanged to independent validation lines.

### Independent validation of the qPCR classifier

The qPCR scoring framework was applied to independent hiPSC lines obtained from external laboratories. These included Test-Q109, Test-Q22, Test-KOLF2_P15, early-passage KOLF2, Test-GTF_IB, Test-CHD_IB and Test-Wt_IB. For each line, RNA extraction, cDNA synthesis, TaqMan qPCR and BCOR-Mut score calculation were performed using the same workflow as for the reference panel.

Where bulk RNA-seq data were available for the same samples, qPCR classifications were compared with transcriptomic clustering and expression of the 15-gene classifier panel.

### Sanger sequencing validation of BCOR exon 7

Genomic DNA was extracted from hiPSC pellets using the Quick-DNA Microprep Kit (Zymo Research). The BCOR exon 7 hotspot region was PCR-amplified using primers flanking the recurrent insertion/deletion site. PCR products were purified and subjected to Sanger sequencing. Sequencing traces were aligned to the BCOR reference sequence using SnapGene software. Indels were manually inspected in electropherograms and compared with the reference exon 7 sequence. In samples classified as BCOR-Mut with no detectable mutation in exon7, other exons are sequenced.

### CRISPR-Cas9 correction of BCOR exon 7 mutation

CRISPR-Cas9 editing was used to correct the recurrent BCOR exon 7 mutation in the BCOR-Mut_93 hiPSC line. The sgRNA was designed to specifically target the mutant BCOR allele by spanning the inserted exon 7 sequence, which is absent from the wildtype allele. A single-stranded homology-directed repair donor oligonucleotide was designed to restore the wildtype BCOR exon 7 sequence and modify the targeted sequence to prevent repeated Cas9 cleavage after successful repair.

Undifferentiated BCOR-Mut_93 hiPSCs were maintained until they reached approximately 70-80% confluency. One hour before nucleofection, the culture medium was replaced with Essential 8 medium supplemented with RevitaCell Supplement and 2 µM M3814/Nedisertib (Selleckchem), a DNA-PKcs inhibitor used to transiently reduce non-homologous end joining and enhance homology-directed repair. Cas9 ribonucleoprotein complexes were assembled by incubating recombinant Cas9 protein with the sgRNA at room temperature according to the manufacturer’s recommendations.

Cells were dissociated into single cells using TrypLE Express Enzyme (Gibco), collected in Essential 8 medium containing RevitaCell Supplement, counted and centrifuged. Approximately 200,000 cells were mixed with the Cas9 ribonucleoprotein complex and the single-stranded donor oligonucleotide, avoiding introduction of air bubbles. The mixture was transferred to a 16-well nucleocuvette strip and electroporated using the SE Cell Line 4D-Nucleofector X Kit S (Lonza) on a 4D-Nucleofector System using program CA137. Cells were rested briefly at room temperature and then plated onto pre-warmed Vitronectin XF-coated plates in Essential 8 medium supplemented with RevitaCell and 2 µM M3814/Nedisertib.

Twenty-four hours after nucleofection, medium was replaced with Essential 8 medium and cells were returned to standard 37°C culture conditions. Forty-eight hours after nucleofection, cells were treated with RevitaCell for 1 h, dissociated to single cells using TrypLE Express, counted and replated at clonal density on Vitronectin XF-coated 100 mm dishes. After approximately one week of culture with daily medium replacement, individual colonies with appropriate hiPSC morphology and round colony edges were mechanically isolated under a stereomicroscope and transferred to Vitronectin-coated 24-well plates.

When colonies reached approximately 80% confluency, each well was split for further expansion and genomic DNA extraction. DNA was extracted using the Quick-DNA Microprep Kit (Zymo Research). The targeted BCOR exon 7 region was PCR-amplified and analysed by Sanger sequencing using SnapGene software. Three precisely corrected clones, BCOR-Res_KI14, BCOR-Res_KI23 and BCOR-Res_KI24, were selected for downstream analyses. A non-rescued edited clone, BCOR-UnRes_KI9, which had undergone the same editing procedure but did not carry the precise correction, was retained as the closest editing-process control.

### UVC irradiation and colony-formation assay

Nucleotide excision repair-deficient hiPSC lines differing in BCOR status were used to assess functional response to UVC-induced DNA damage. hiPSCs were cultured in replicate plates until they reached approximately 80-90% confluency. For irradiated plates, Essential 8 medium was removed and cells were washed twice with 1x DPBS. Cells were covered with room-temperature DPBS and exposed to a single UVC dose of 0.75 J/m2 in a UVC irradiation chamber. After irradiation, DPBS was replaced with Essential 8 medium supplemented with Penicillin-Streptomycin and cells were returned to a 37°C incubator. Matched non-irradiated plates were processed in parallel.

Twenty-four hours after irradiation, irradiated and non-irradiated cells were dissociated to single cells using TrypLE Express Enzyme, washed, counted and replated at 20,000 cells per well in Vitronectin XF-coated 6-well plates. Cells were cultured in Essential 8 medium supplemented with 1x RevitaCell Supplement, with daily medium changes, until control cultures reached confluency. Cells were then washed with DPBS and stained with 0.5% crystal violet prepared in 25% methanol for 30 min at room temperature. Plates were rinsed several times with double-distilled water, inverted and left to dry for 48 h before imaging.

### Statistical analysis and data visualisation

All statistical analyses and visualisations were performed in R. RNA-seq differential expression was analysed using DESeq2, with adjusted P value < 0.05 considered significant. Proteomic differential abundance was analysed using Student’s t tests in Perseus with the thresholds described above. Fisher’s exact tests were used for categorical enrichment analyses, including exon-level mutation enrichment and pathway enrichment. Pearson correlations were used to assess concordance between RNA-seq, proteomic and chromatin datasets.

Principal component analysis, volcano plots, heatmaps, bar plots, correlation plots and qPCR classifier plots were generated using R packages including ggplot2, pheatmap, ComplexHeatmap, EnhancedVolcano, and tidyverse packages. ChIP-seq heatmaps and profile plots were generated using deepTools. Final figures were assembled in Adobe Illustrator.

## Data and code availability

This paper did not generate any original code. BCOR somatic mutation calling data is available at: https://doi.org/10.1038/s41588-022-01147-3 and https://doi.org/10.1016/j.xgen.2023.100280. RNAseq data of hiPSCs can be accessed here: https://doi.org/10.1038/s41588-022-01147-3 and KDM2B-KO RNAseq data generated by Wang. Z, *et al*. can be accessed here: https://doi.org/10.1016/j.stem.2017.12.002. ChIPseq and proteomics datasets will be made publicly available upon acceptance of this paper.

BsCORe website is now accessible for editors and reviewers using an ID and password and will be publicly available upon acceptance of the paper.

---

**Supplementary Figure 1.**
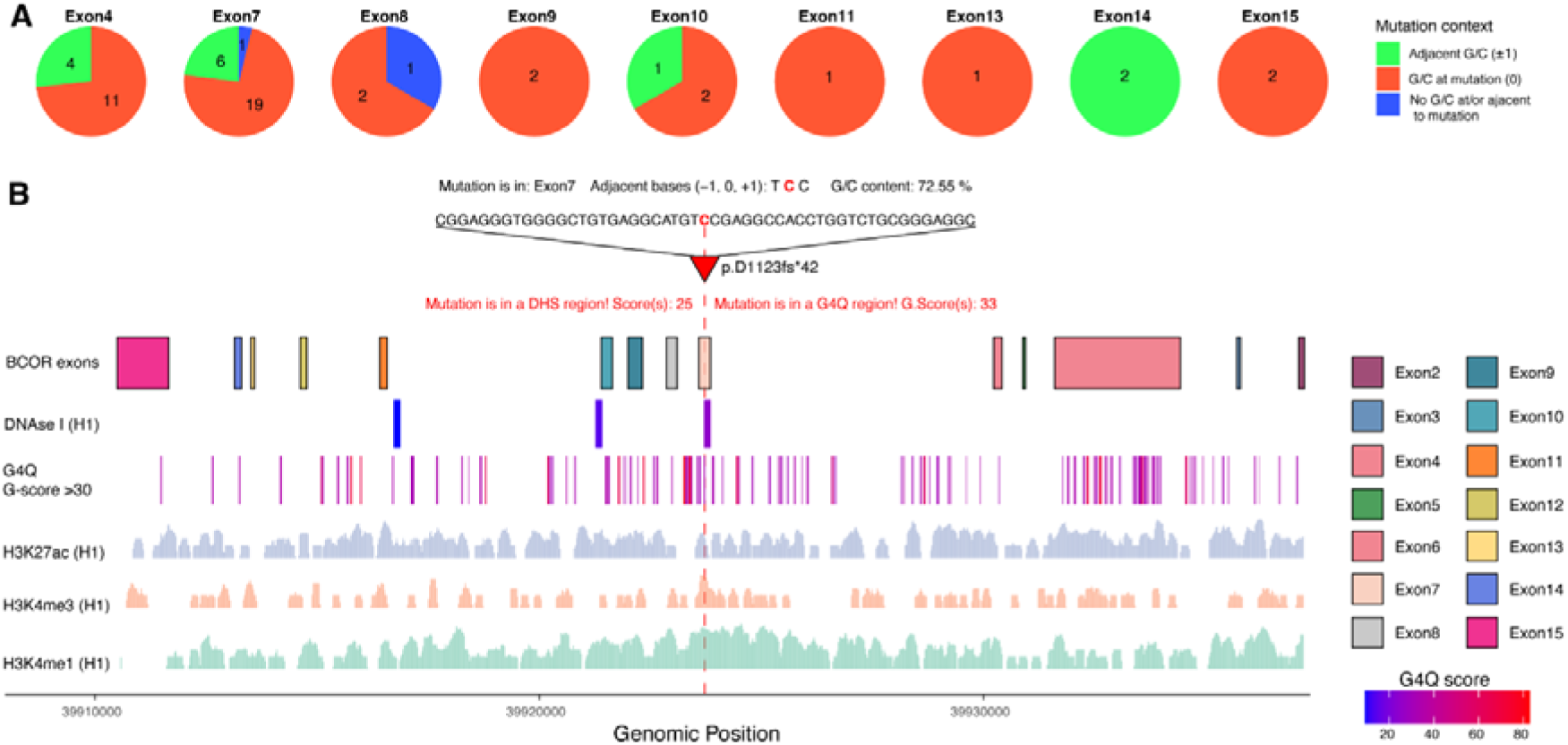
Recurrent BCOR mutations in hiPSCs occur within GC-rich sequence contexts. A, Sequence context of BCOR mutations identified in hiPSCs stratified by exon. Mutations were classified according to whether the altered nucleotide occurred at a G/C base, adjacent to a G/C base (±1 bp), or outside a local G/C-rich context. **B,** Genomic landscape surrounding a recurrent BCOR exon 7 hotspot centred on p.D1123fs*42. Tracks show BCOR exon structure, DNase I hypersensitivity sites, predicted G-quadruplexes (G4Q; G-score >30), and chromatin profiles.

**Supplementary Figure 2.**
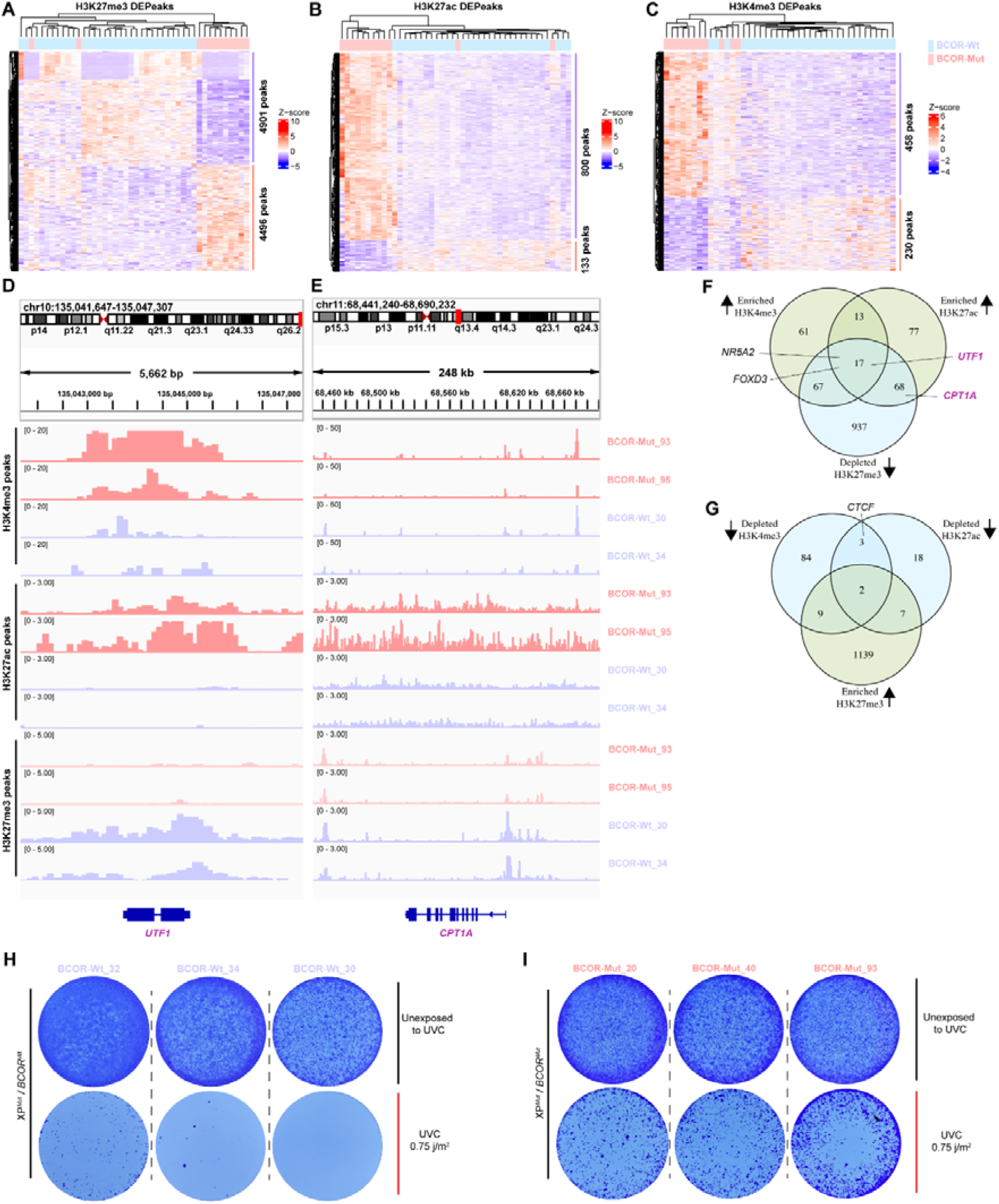
BCOR mutations induce genome-wide chromatin alterations and increased tolerance to DNA damage. A–C, Heatmaps of differential histone modification enrichment between BCOR-Wt and BCOR-Mut hiPSCs (padj < 0.05) for H3K27me3 (**A**), H3K27ac (**B**) and H3K4me3 (**C**). Heatmaps display row-scaled normalized signal intensities (Z-scores). **D,E,** Representative genomic loci showing coordinated chromatin remodelling in BCOR-Mut hiPSCs at the *UTF1* (**D**), and *CPT1A* (**E**) loci. Tracks are shown for two BCOR-Mut lines (BCOR-Mut_93 and BCOR-Mut_95) and two BCOR-Wt lines (BCOR-Wt _30 and BCOR-Wt_34). **F,** Overlap of genes associated with increased H3K4me3 and H3K27ac enrichment together with depletion of H3K27me3. **G,** Overlap of genes associated with depletion of H3K4me3 and H3K27ac and enrichment of H3K27me3. **H,I,** Colony formation assays following UVC irradiation (0.75 J m_⁻_²) in nucleotide excision repair-deficient XP hiPSCs with BCOR-Wt (**H**) or BCOR-Mut (**I**) backgrounds.

**Supplementary Figure 3.**
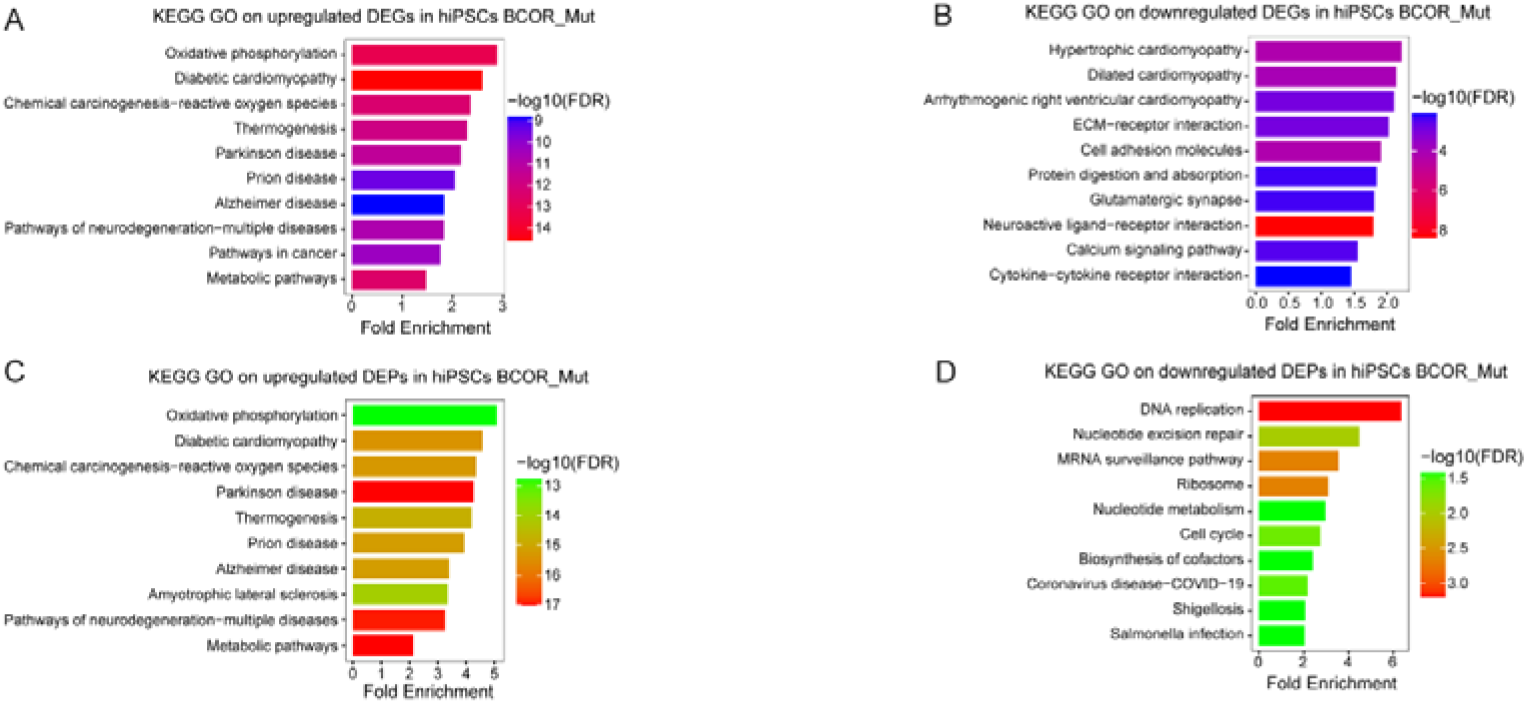
Pathway enrichment analyses reveal upregulation of metabolic signalling-associated programmes in BCOR-mutant hiPSCs. A,B, KEGG pathway enrichment analyses of DEGs identified by RNA-seq. Enriched pathways among upregulated genes (**A**) and downregulated genes (**B**) in BCOR-Mut relative to BCOR-Wt B-hiPSCs. **C,D,** KEGG pathway enrichment analyses of DEPs identified by proteomics. Enriched pathways among proteins increased (**C**) and decreased (**D**) in BCOR-Mut B-hiPSCs relative to BCOR-Wt B-hiPSCs.

**Supplementary Figure 4.**
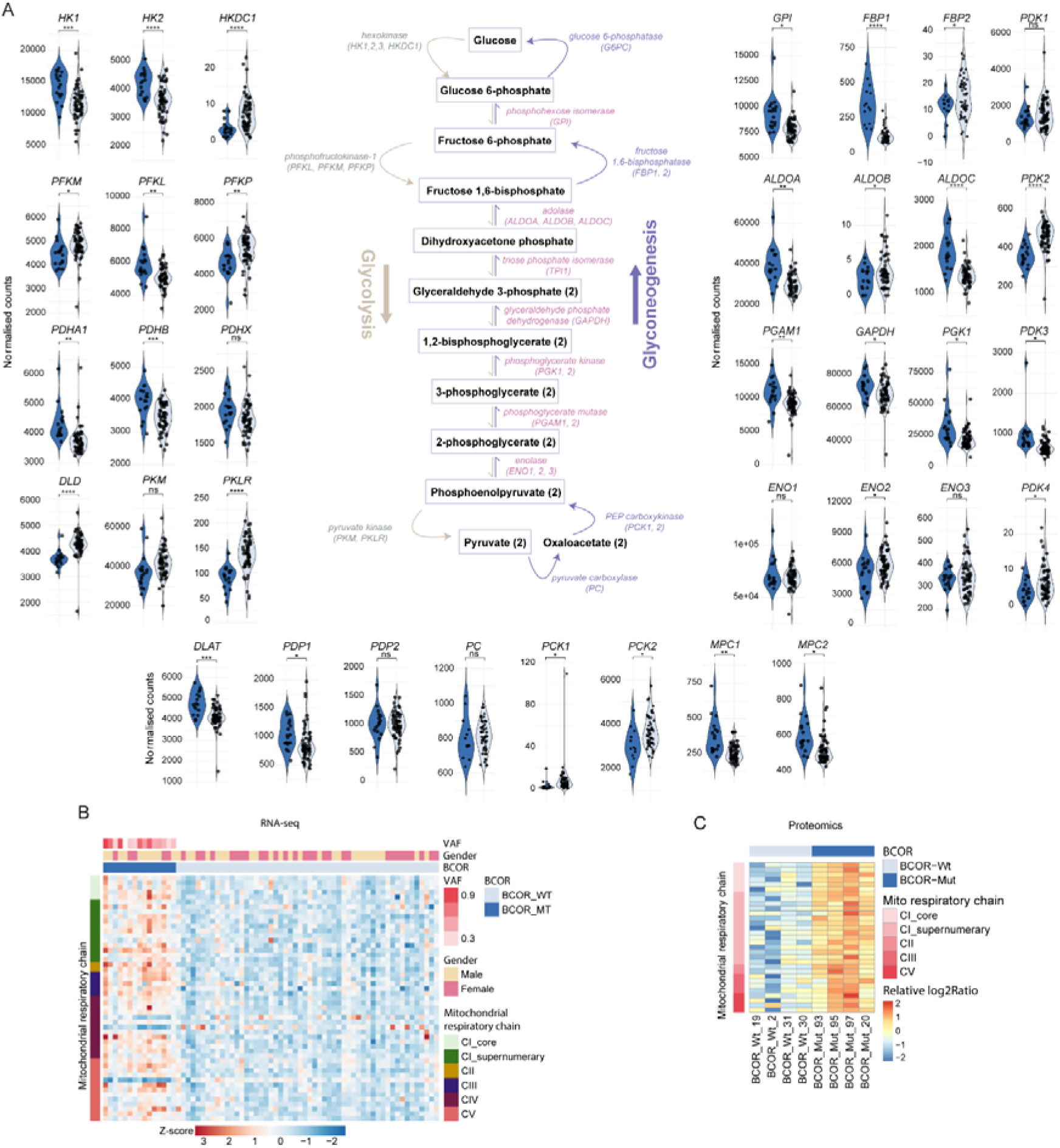
BCOR-mutant hiPSCs show a dysregulated glucose metabolism and coordinated activation of mitochondrial respiratory chain. **A**, Expression of genes involved in glucose metabolism in BCOR-mutant and BCOR-wildtype hiPSC. **B**,**C**, Heatmaps showing expression of mitochondrial respiratory chain components identified by RNA-seq (B) and proteomics (C). Statistical significance was assessed using a two-sided t-test (*p* < 0.0001****, *p* < 0.001***, *p* < 0.01**, *p* < 0.05*, ns = non-significant).

**Supplementary Figure 5.**
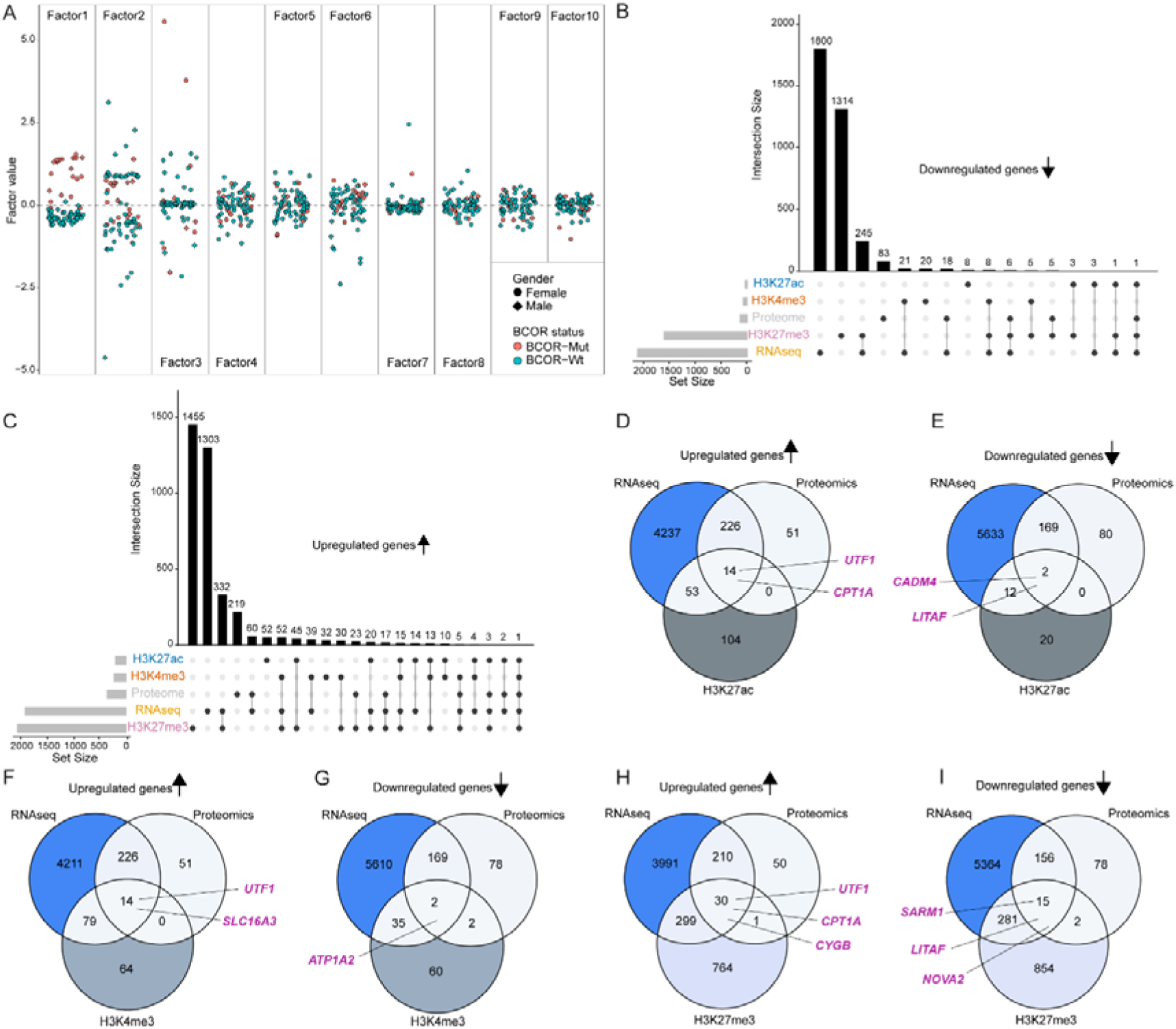
BCOR-associated molecular signature across chromatin, transcriptomic and proteomic datasets. A, Distribution of samples across the first ten latent factors identified by multi-omics factor analysis (MOFA). Factor 1 showed the strongest separation between BCOR-Wt and BCOR-Mut samples. **B,C,** UpSet plots showing overlap of differential features identified across RNA-seq, proteomic, H3K27ac, H3K27me3 and H3K4me3 datasets. Downregulated features (B) and upregulated features (C). **D,E,** Overlap between DEGs, DEPs and genes associated with differential H3K27ac regions. **F,G,** Overlap between DEGs, DEPs and genes associated with differential H3K4me3 regions. **H–I,** Overlap between DEGs, DEPs and genes associated with differential H3K27me3 regions.

**Supplementary Figure 6.**
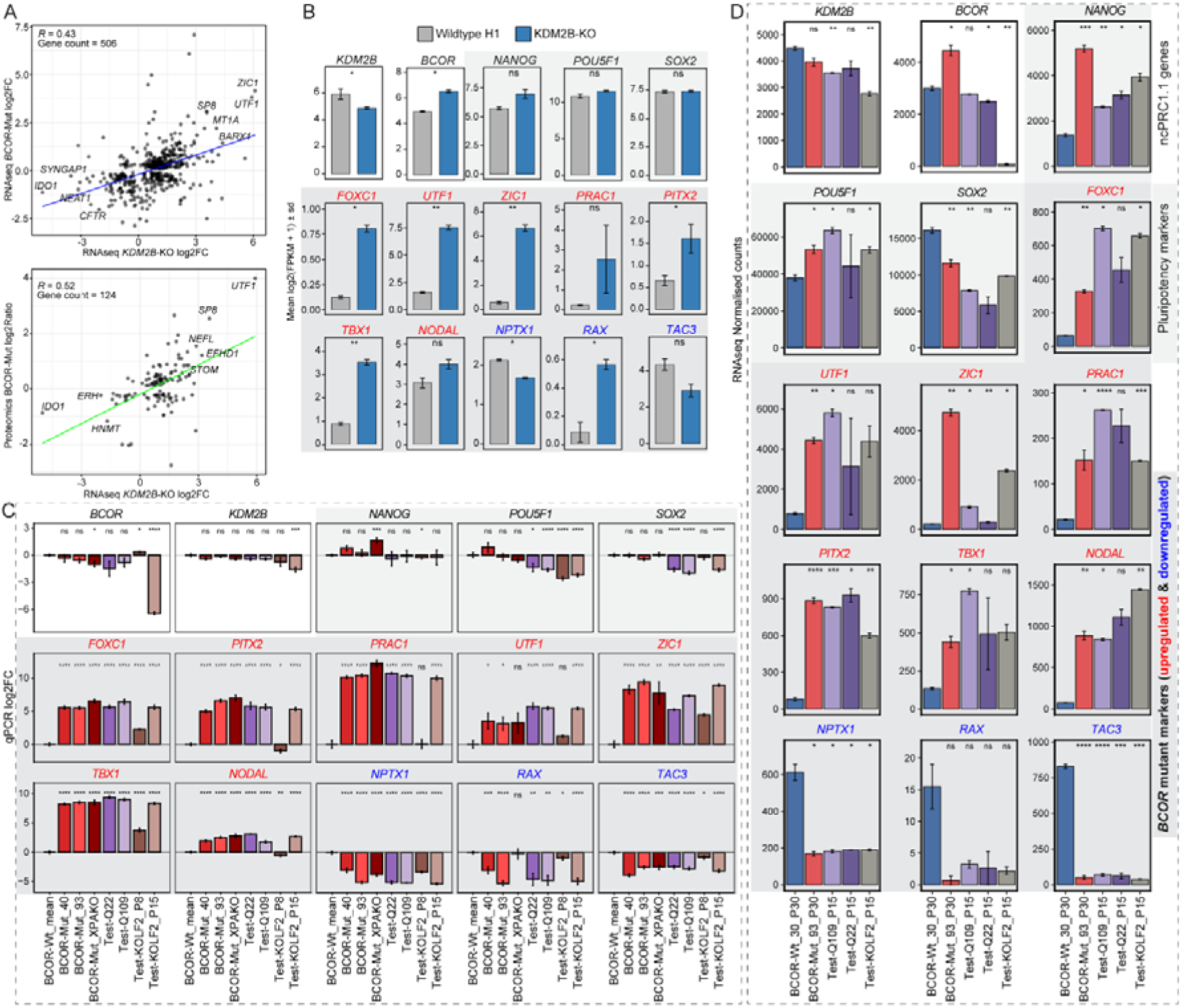
Development of a qPCR classifier and identification of BCOR-mutant hiPSC lines. **A**, Correlation of transcriptomic changes between BCOR-mutant hiPSCs and an independent KDM2B-knockout hESCs dataset. **B**, Expression of BCOR-mutant markers in wild-type H1 and KDM2B-knockout cells (data from Wang et al., 2018, Cell Stem Cell 22, 235–251). Values are shown as mean log_₂_(FPKM + 1) ± s.d. **C**, qPCR analysis of reference BCOR-mutant lines and independent test samples. Expression values are shown as log_₂_ fold change relative to the mean BCOR-Wt reference (mean of BCOR-Wt_30, BCOR-Wt_33, BCOR-Wt_34) **D**, RNA-seq validation of ncPRC1.1 genes, pluripotency markers and BCOR-mutant markers. Values shown are relative to BCOR-Wt_30_P30. Statistical significance was assessed using a two-sided t-test (*p* < 0.0001****, *p* < 0.001***, *p* < 0.01**, *p* < 0.05*, ns = non-significant).

